# Spatiotemporal Clusters of ERK Activity Coordinate Cytokine-induced Inflammatory Responses in Human Airway Epithelial Cells

**DOI:** 10.1101/2024.02.03.578773

**Authors:** Nicholaus L. DeCuzzi, Daniel P. Oberbauer, Kenneth J. Chmiel, Michael Pargett, Justa M. Ferguson, Devan Murphy, Amir A. Zeki, John G. Albeck

## Abstract

**RATIONALE:** Spatially coordinated ERK signaling events (“SPREADs”) transmit radially from a central point to adjacent cells via secreted ligands for EGFR and other receptors. SPREADs maintain homeostasis in non-pulmonary epithelia, but it is unknown whether they play a role in the airway epithelium or are dysregulated in inflammatory disease.

**OBJECTIVES:** (1) To characterize spatiotemporal ERK activity in response to pro-inflammatory ligands, and (2) to assess pharmacological and metabolic regulation of cytokine-mediated SPREADs.

**METHODS:** SPREADs were measured by live-cell ERK biosensors in human bronchial epithelial cell lines (HBE1 and 16HBE) and primary human bronchial epithelial (pHBE) cells, in both submerged and biphasic Air-Liquid Interface (ALI) culture conditions (i.e., differentiated cells). Cells were exposed to pro-inflammatory cytokines relevant to asthma and chronic obstructive pulmonary disease (COPD), and to pharmacological treatments (gefitinib, tocilizumab, hydrocortisone) and metabolic modulators (insulin, 2-deoxyglucose) to probe the airway epithelial mechanisms of SPREADs. Phospho-STAT3 immunofluorescence was used to measure localized inflammatory responses to IL-6.

**RESULTS:** Pro-inflammatory cytokines significantly increased the frequency of SPREADs. Notably, differentiated pHBE cells display increased SPREAD frequency that coincides with airway epithelial barrier breakdown. SPREADs correlate with IL-6 peptide secretion and localized pSTAT3. Hydrocortisone, inhibitors of receptor signaling, and suppression of metabolic function decreased SPREAD occurrence.

**CONCLUSIONS:** Pro-inflammatory cytokines modulate SPREADs in human airway epithelial cells via both secreted EGFR and IL6R ligands. SPREADs correlate with changes in epithelial barrier permeability, implying a role for spatiotemporal ERK signaling in barrier homeostasis and dysfunction during inflammation. The involvement of SPREADs in airway inflammation suggests a novel signaling mechanism that could be exploited clinically to supplement corticosteroid treatment for asthma and COPD.

**Brief Summary:** Combining live-cell ERK biosensors with multiple human airway epithelial models, we demonstrate that pro-inflammatory cytokines cause spatiotemporally organized ERK signaling events called “SPREADs”, correlating with conditions that disrupt epithelial barrier function. Additionally, common anti-inflammatory treatments tocilizumab, gefitinib, and hydrocortisone suppress cytokine-induced SPREADs. These findings suggest that localized ERK signaling coordinates the innate immune response via spatially restricted cytokine release and regulation of airway barrier permeability.

## INTRODUCTION

Epithelial cells line the mammalian airway and protect the lung from harmful inhaled particles and pathogens. Such insults trigger inflammatory cytokine signaling within the context of respiratory diseases, as exemplified by asthma and chronic obstructive pulmonary disease (COPD) (1–9). In these diseases, signaling within the epithelial layer is dysregulated at a broad level, but how do the behaviors of individual cells contribute to this disruption? Within an epithelial surface, cells can respond independently to cytokines, but multicellular, coordinated cascades of responses serve to efficiently propagate stimuli (10). Such collective, timed behaviors may be essential to elicit innate immune defenses and maintain tissue barrier homeostasis within the airway epithelium.

Extracellular signal-regulated kinase (ERK) is a well-established regulator of epithelial cell behavior with a plurality of functions in wound healing, tobacco smoke reactivity, inflammation, and innate immune responses. ERK plays a role in these behaviors through its effects on cell proliferation, death, epithelial-mesenchymal transition (EMT), and cytokine production (5, 11, 12). Also implicated in previous studies is the integrity of the epithelial barrier itself, which under influence of ERK signaling allows neutrophil influx from sub-epithelial regions across an intact epithelium (13). Given this widespread role in airway disease pathogenesis and the underlying mechanisms, targeting the ERK 1/2 pathway may have therapeutic potential (14, 15).

In many tissues, ERK signaling is highly localized and temporally dynamic (16), but such patterns have not been investigated in the airway. Given ERK’s known roles in the airway epithelium, a possible function for spatially localized ERK signaling is the coordination of cytokine production with epithelial barrier permeability, creating focal points for immune cell entry. Paracrine release of EGFR ligands often causes a unique type of ERK activity localized to regions of 5 to 50 cells (17–19). These spatially localized ERK signaling events have been termed “SPREADs”, for Spatial Propagation of Radial ERK Activity Distributions, and were first described in non-human epithelial cells (17, 18). Since then, SPREADs have been linked to cell fate events in various non-pulmonary epithelial systems, including instances of proliferation (18), cancer cell ejection (20), and apoptosis (19). However, the role of SPREADs in the context of the airway and their modulation by proinflammatory conditions remains unknown. We hypothesized a new function for SPREADs in airway epithelial cells, in which they allow inflammatory stimuli to transiently increase barrier permeability and cytokine production during airway inflammation.

To evaluate the potential role of SPREADs in airway inflammation, several questions must be addressed. First, while secreted EGFR ligands have been implicated in the propagation of ERK activity, it is unknown if secreted pro-inflammatory cytokines such as IL-6 are connected to SPREAD behavior. Second, it will be important to determine if SPREADs are affected by the metabolic environment of the epithelium, as patients with metabolic disorders (obesity, diabetes) are more susceptible to lung inflammation. Finally, determining the effects of known anti-inflammatory treatments such as corticosteroids on SPREAD behavior is needed to establish the clinical relevance of spatial ERK regulation.

To address these gaps in knowledge, we integrated fluorescent biosensors for ERK activity into human airway epithelial cells and collected live-cell ERK activity data in response to pro-inflammatory conditions, with single cell resolution. We demonstrate that SPREADs occur in various airway epithelial cell lines and primary cells, co-localize with secondary IL-6 secretion, and are modulated by metabolic factors and anti-inflammatory drugs. Our findings suggest a spatiotemporally dynamic mechanism underlying airway disease progression and provide insight into the effects of pro-inflammatory cytokines on airway epithelial cell signaling

## METHODS

### Human Airway Epithelial Cell Sources

HBE1, a papilloma virus-immortalized human bronchial epithelial cell line originally generated by Dr. James Yankaskas (University of North Carolina, Chapel Hill, NC) (21), was provided by Dr. Reen Wu of University of California, Davis, and was cultured as described in (22). 16HBE14o-(referred to as 16HBE) cells were obtained from Sigma-Aldrich (#SCC150) and cultured in ɑ-MEM (Sigma M2279) + 2mM L-Glutamine + 10% FBS (23). Normal human primary tracheobronchial epithelial cells (pHBE) were isolated from human bronchi and distal trachea obtained from organ donors via the Cooperative Human Tissue Network. pHBE cells were cultured in submerged and air-liquid interface (ALI) culture as described in (24). *See supplemental methods for further details*.

### Live Cell Measurement of ERK activity

**Figure 1A** shows example images of the ERK biosensors ERK-KTR (25), which has a typical dynamic range of 2-2.5 (arbitrary units) and EKAREN4 (26), which has a typical dynamic range of 0.2-0.25 (arbitrary units); the biosensors are further described in (27). ERK-KTR or EKAREN4 biosensors were stably integrated into cells by lentiviral transduction and used to record single-cell ERK activity at 6-minute intervals for 24 hours following treatment (**Sup. Video 1**). For live-cell imaging, cells were deprived of EGF and BNE/FBS for at least 6 hours (following the protocol described in (28)) and then exposed to one of six conditions that were designed to recapitulate various airway inflammatory states (**Table 1**), individual pro-inflammatory cytokines from these conditions, or controls. In certain experiments, cells were fixed and immunofluorescence staining was used to measure pSTAT3 (**Figure 1B**). Single-cell ERK activity was quantified and pulse analysis was performed using MATLAB as described in (29). *See supplemental methods for further details*.

**Figure 1.**
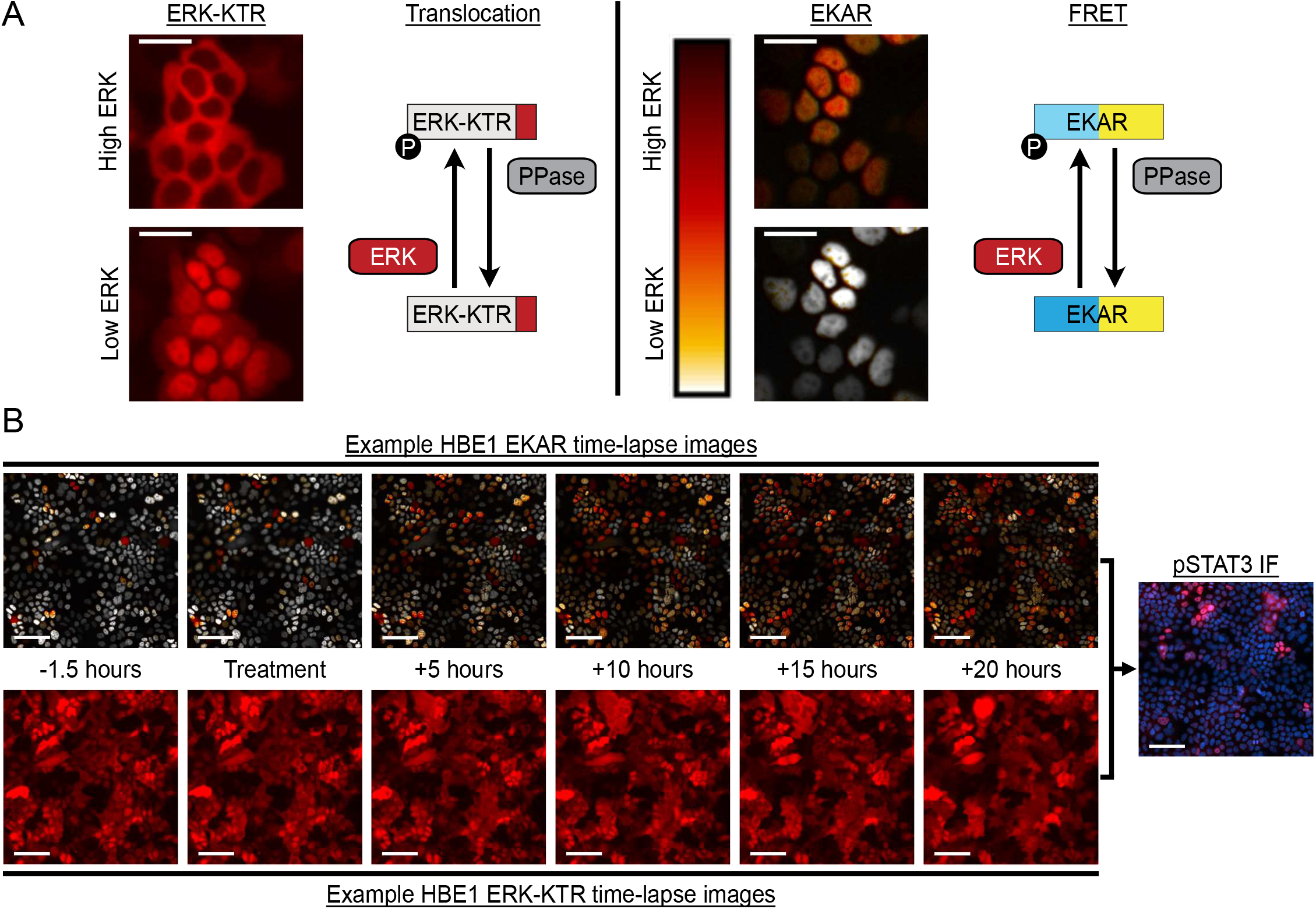
ERK biosensor diagrams, and example images. **(A)** Example images of HBE1 cells expressing fluorescent biosensors for ERK activity, with the top image showing near-maximal ERK activity (EGF stimulation) and the bottom image showing near-minimal activity (MEK inhibitor). *Left*, ERK-KTR (translocation reporter) localizes to the cytoplasm upon ERK activation and to the nucleus upon ERK inhibition. *Right*, EKAREN4 (FRET reporter) shows a high FRET signal upon ERK activation and a low FRET signal upon inhibition. Scale bars, 25 μm. **(B)** Example time lapse images of HBE1 cells expressing both the ERK-KTR and EKAREN4 biosensors. *Right panel:* for certain experiments, cells were fixed following live-cell imaging of the biosensor and stained for pSTAT3 (red) and DNA (blue). Scale bars, 100 μm.

**Table 1:**
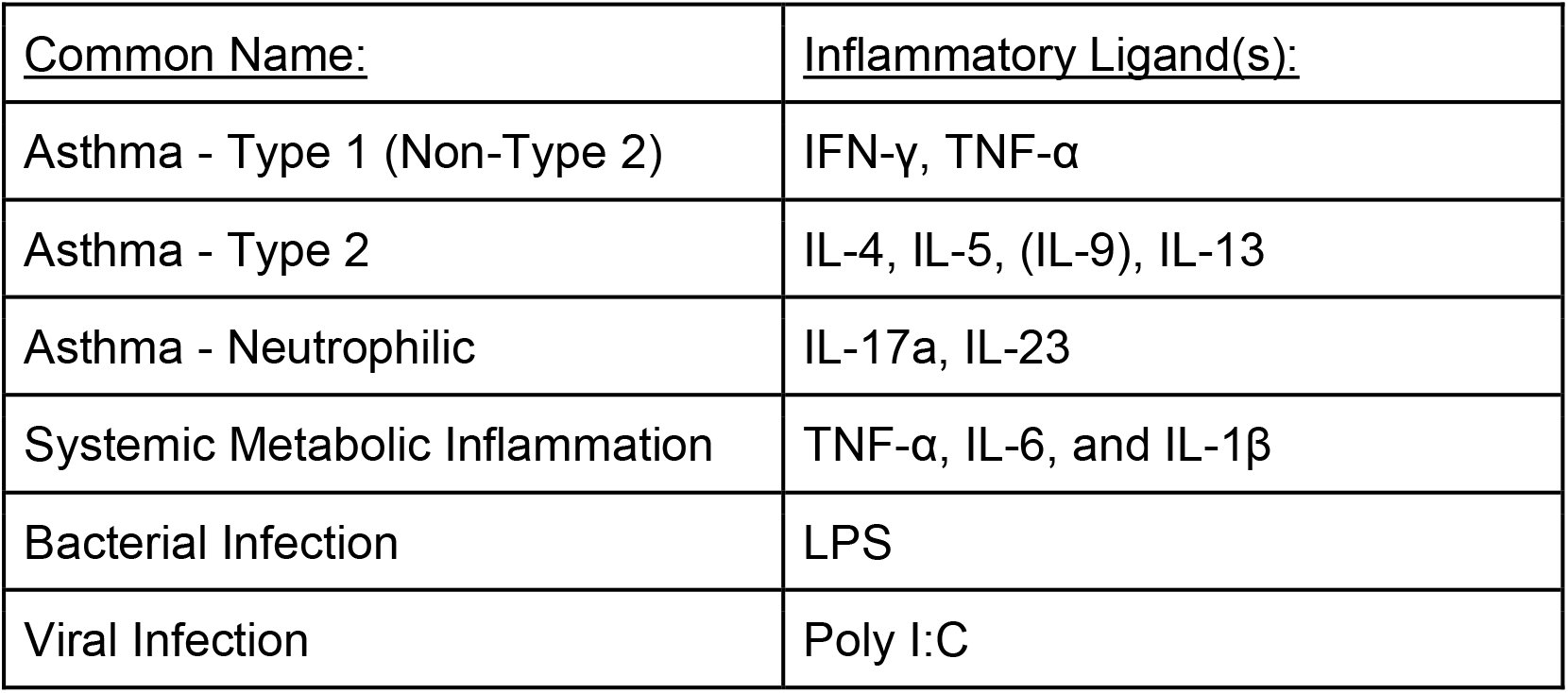
Groups of pro-inflammatory conditions tested.

### Quantification of Spatially Localized ERK Activity

We developed a MATLAB implementation of Automated Recognition of Collective Signaling (ARCOS) (19, 30). Following the ARCOS methodology, ERK pulses detected in time series data were binarized and subsequently clustered with Density Based Spatial Clustering of Applications with Noise (DBSCAN). Cluster labels were tracked using the k-nearest neighbors algorithm to match neighbors of clusters within a given distance between time points (epsilon).

Our implementation expands on ARCOS by using pulse analysis to identify active ERK pulses for binarization, adding automatic estimations of epsilon and the minimum number of points within epsilon distance of a point for that point to be considered a core point, based on the distributions of the input data and cluster area calculations using MATLAB’s native boundary tracing method. Cluster data can be filtered by duration, start time, maximum area, maximum count, mean rate of change of count and mean rate of change of area, effectively narrowing down the pool of spread candidates. We validated our approach by generating synthetic spreads and using Adjusted Rand Indices (ARI) to compare per-timepoint ground truth cluster assignments to ARCOS cluster assignments of varying lifetime and frequency, observing mean per-timepoint ARI values between 0.96 and 0.99. (Code and methods available at our github repository: https://github.com/Albeck-Lab/ARCOS-MATLAB) *See supplemental methods for further details*.

### Trans-Epithelial Electrical Resistance (TEER) Measurement

TEER values were measured in Phosphate-Buffered Saline (with Ca^2+^ / Mg^2+^) in air-liquid-interface (ALI inserts by Corning, 3460) using a Trans-Epithelial Electrical Resistance Meter (EVOM3 -World Precision Instruments), which was used and optimized according to the manufacturer instructions.

### Measurement of Secreted Ligands

Cultured HBE1 cells were immersed in serum-free media and treated with cytokines IL-1β (10 ng/ml) and IL-6 (10 ng/ml) with media collections at 2- and 12-hour timepoints. Cytokine levels in media, in treated and untreated wells, were measured via a Luminex High Performance custom assay plate (MilliporeSigma, Item no. HCYTOMAG-60K-16) for detection of TNF-α, RANTES, MCP-1, MIP-1α, Eotaxin, and VEGF-A interleukins 1α, 1β, 4, 6, 8, 10, 17A, 17E, 17F, and 18. Cytokine array measurements were made using the Proteome Profiler Human XL Cytokine Array Kit (ARY022B) from R&D systems, following the manufacturer’s instructions.

## RESULTS

### Airway disease conditions induce dynamic ERK activity in HBE1 cells

To evaluate temporal ERK activity modulated by pro-inflammatory cytokines, we generated HBE1, 16HBE, and primary human bronchial epithelial (pHBE) cells expressing ERK biosensors (**Figure 1, Sup. Video 1**). For live-cell imaging, cells were deprived of EGF and BNE/FBS for at least 6 hours and then exposed to one of six conditions that were designed to recapitulate various airway inflammatory states (**Table 1**), individual cytokines from these conditions, or controls.

Across the treatments, we observed differences in the fraction of cells responding and in the intensity and duration of those responses. While the vehicle control induced a minimal response (**Figure 2A**), the canonical growth factor EGF induced a maximal ERK response, which lasted 60 to 120 minutes, with 99% of cells responding to the treatment (**Figure 2B, Figure E1A**). After this initial period, mean ERK activity tapered to approximately half-maximal levels for the remainder of the experiment (22 to 24 hours) (**Figure 2B**). The combination of IL-1β, IL-6, and TNF-α induced a strong immediate ERK response, which on average lasted 24 minutes with 76% cells responding (**Figure 2C, Figure E1A**). When tested individually, IL-1β or IL-6 treatment elicited initial ERK responses lasting 19 minutes or 21 minutes, and 61% and 51% of cells responding, respectively (**Figure 2D&E, Figure E1A**). While 13% of cells responded to IFN-γ and TNF-α (**Figure 2F, Figure E1A**), responder activity also lasted 24 minutes. Most other pro-inflammatory condition groups tested (IL-4/5/9/13, IL-17A/23, or LPS) did not elicit an initial ERK response that was significantly different from the vehicle (negative control) treatment (7% responders, **Figure E1A, Figure E2A-C**); however, poly I:C treatment induced a delayed initial peak that was variable across experiments (**Figure E2D&E**).

**Figure 2.**
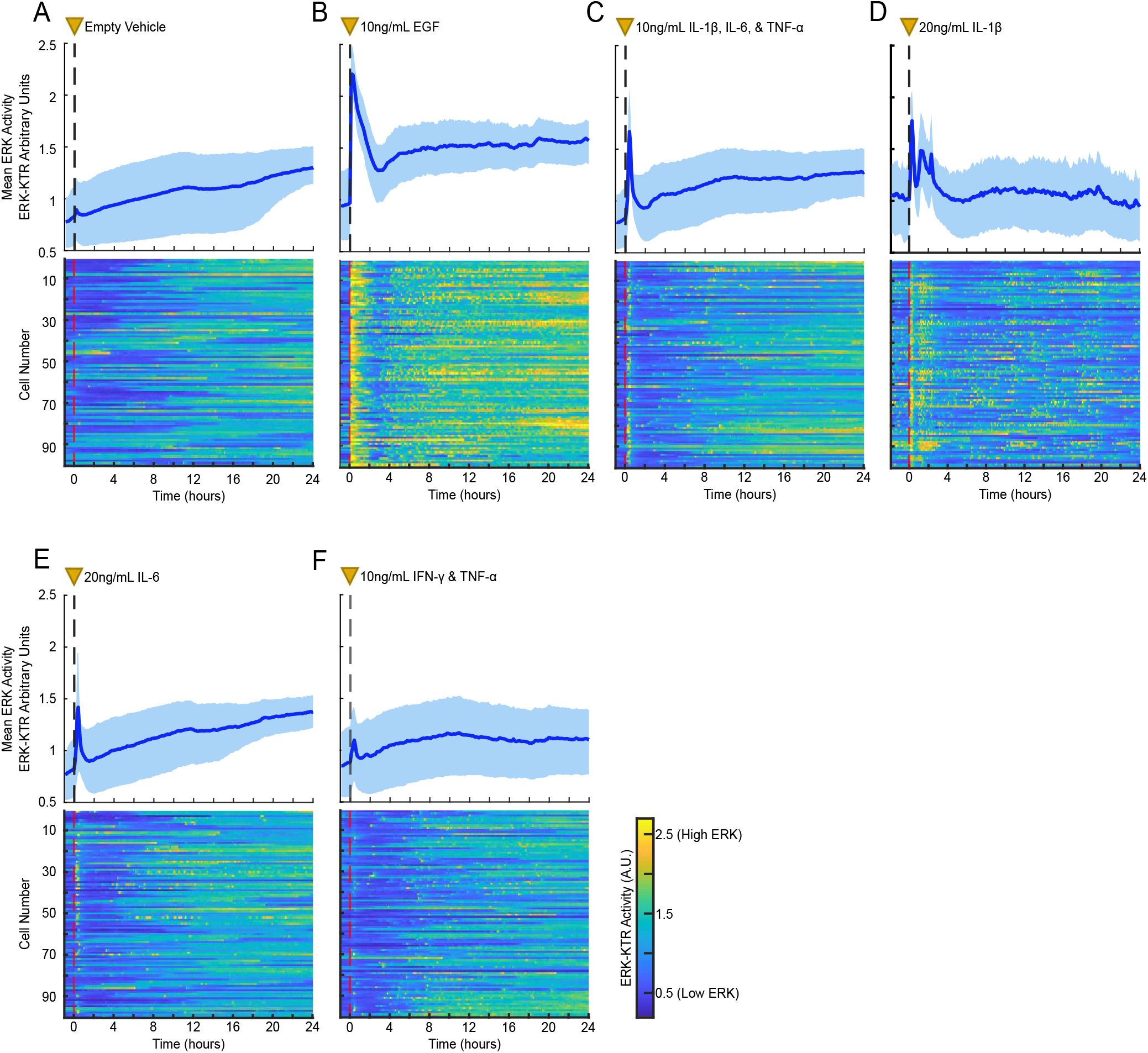
ERK activity of HBE1 cells in response to vehicle control, EGF, or pro-inflammatory ligands. **(A-F)** The top panels show the mean ERK-KTR signal (dark blue line) with 25th/75th quantiles (shaded light blue regions). The bottom panels show heatmaps of ERK activity (color axis) for 100 randomly selected cells. Each horizontal row depicts one cell’s activity over time. Heatmaps show representative data from 1 of 3 technical replicates, >500 cells per condition. The dashed vertical lines and yellow triangles (0 hours on the time axis) indicate when treatments (listed next to the yellow triangles) were applied to the cells.

At the single-cell level, we observed significant heterogeneity in long-term ERK activity profiles under pro-inflammatory cytokines (**Figure 2 - heatmaps, Figure E1B**). While vehicle-treated cells displayed only rare sporadic ERK activation (0.10 ERK pulses per hour), EGF-treated cells showed greatly increased mean activity and frequency of ERK pulses (0.41 pulses per hour). Notably, HBE1 cells treated with IL-1β/IL-6/TNF-α, IL-1β, IL-6, TNF-α, IFN-γ/TNF-α, or IFN-γ displayed increased sporadic ERK activation (0.13 to 0.27 pulses per hour), but without strongly increasing the mean ERK activity (**Figure 2 & Figure E2B**). Excluding the initial treatment response, we found no statistical differences in pulse duration in cytokine treated cells compared to vehicle control (mean pulse duration = 16 minutes; **Figure E1C**).

### Pro-Inflammatory cytokines modify SPREADs in airway epithelial cell lines

We next evaluated if the sporadic ERK pulses induced by pro-inflammatory cytokines were spatiotemporally localized. Concentric ERK signaling waves (i.e. SPREADs) were clearly visible in time-lapse images (e.g., **Figure 3A, Sup. Video 2**). To evaluate whether SPREAD behavior is modified by pro-inflammatory cytokines, we used a SPREAD detection program (ARCOS; **Figure 3B**) to analyze SPREAD features including frequency of occurrence (SPREADs per hour normalized to mm^2^), duration, maximum area, and localization of SPREAD events.

**Figure 3:**
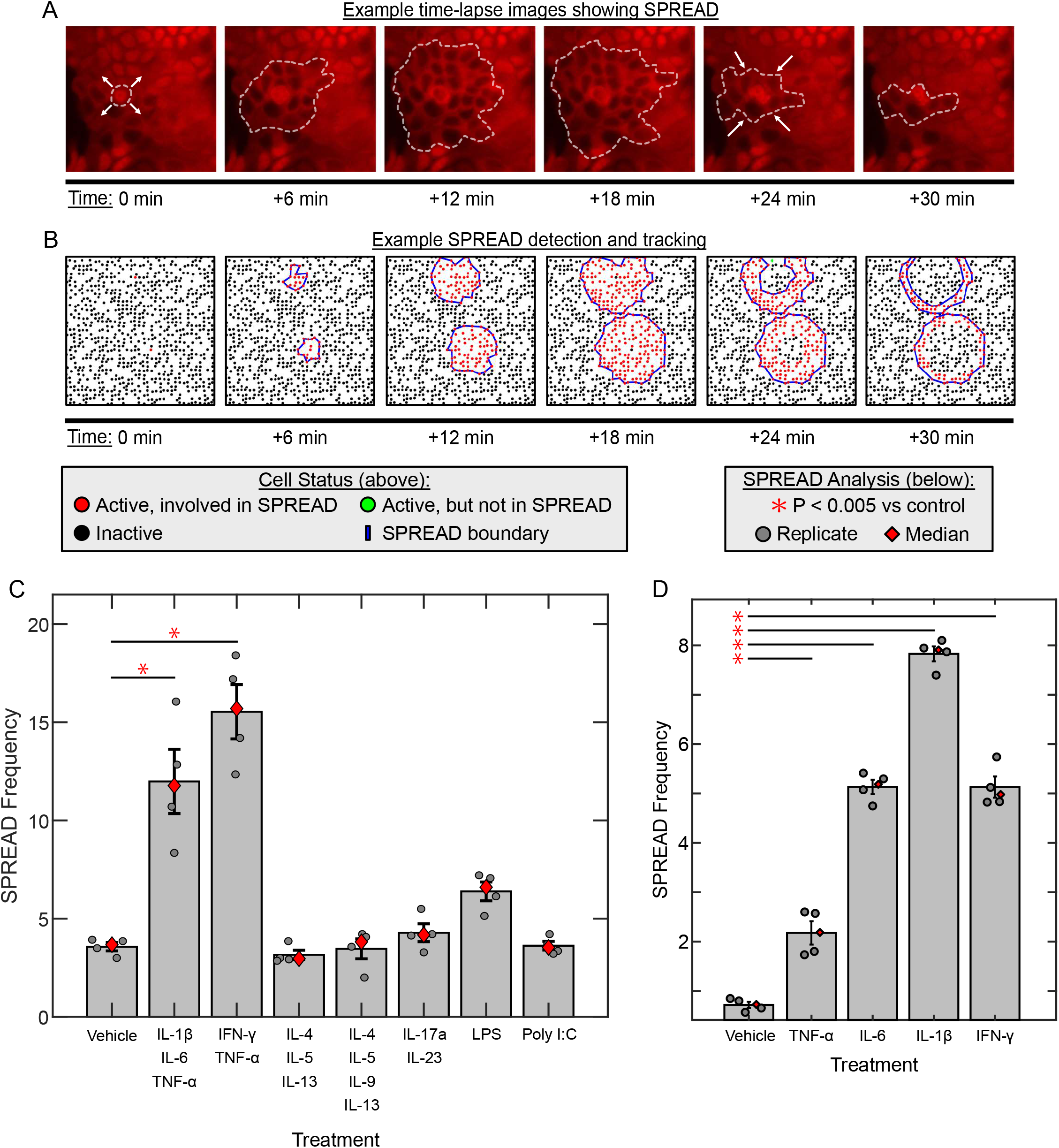
Spatially localized ERK activity in HBE1 cells in response to pro-inflammatory ligands. **(A)** Time lapse images showing an example ERK activity wave (SPREAD) occurring in HBE1 cells. The dashed white line indicates the edge of a SPREAD event, and the white arrows represent the direction of SPREAD growth or shrinkage. Time 0 indicates the initiation point of the example SPREAD, approximately 16 hours after treatment with 10 ng/mL IL-1β. **(B)** SPREAD detection results on simulated ERK activity data. Red dots represent active cells identified to be part of a SPREAD. Blue solid lines show the boundary of detected SPREADs. Black dots represent inactive cells. Green dots show cells that are considered to have active ERK, but are not a part of a detected SPREAD. **(C and D)** SPREAD frequency (SPREAD occurrences per hour, normalized to mm^2^) in the 24 hours following treatment with the indicated stimuli. Gray dots represent technical replicates, red diamonds show the median, and error bars show S.E.M. Red asterisks and lines indicate conditions where SPREAD frequency is significantly different compared to the control group (P < 0.005), calculated using 1-way ANOVA compared to the control and adjusted for multiple comparisons via the Dunnett’s procedure.

HBE1 cells displayed a significant increase in SPREAD frequency, compared to vehicle-treated cells, only when treated with combinations of either IL-1β/IL-6/TNF-α or IFN-γ/TNF-α (**Figure 3C**). To determine which cytokines in the bulk treatment groups caused the increased SPREAD frequency, we tested each cytokine separately. TNF-α, IL-6, IL-1β, or IFN-γ (20 ng/mL) all caused a significant increase in SPREAD frequency compared to their vehicle-treated counterparts (**Figure 3D**). In EGF-treated cells, the detected SPREAD frequency was inconsistent across replicates and thus was excluded from analysis (**Figure E3A; Sup. Video 3**).

In addition to increased frequency of SPREADs, the average area of SPREADs were significantly larger in IL-1β/IL-6/TNF-α and IFN-γ/TNF-α treated cells (1600μm^2^ and 1500μm^2^) compared to vehicle control (600μm^2^) (**Figure E3B**). However, despite heterogeneity in the duration of SPREADs within each condition, no condition displayed a significantly different mean SPREAD duration compared to control conditions (means ranging from 20 minutes to 25 minutes, with a 22 minute average, **Figure E3C**).

To evaluate SPREADs in an alternate epithelial model, we performed a similar analysis in 16HBE cells and observed both common features in SPREAD behavior as well as notable differences. As with HBE1 cells, 16HBE responded with an initial peak of ERK activity when stimulated by EGF, IL-1β/IL-6/TNF-α, or IL-1β stimulation (**Figure 4A-D**). However, 16HBE cells did not display a strong ERK response to IL-6 or IFN-γ/TNF-α (**Figure 4E & F**). A delayed response to Poly I:C, beginning 1 hour after treatment was also present (**Figure 4G**). Also consistent with HBE1 cells, 16HBE cells did not respond to IL-4/5/9/13, IL-17A/23, or LPS (**Figure E4**). At the single-cell level, individual 16HBE cells displayed sporadic peaks of ERK activity over extended treatment times, and many of these peaks were organized as SPREADs (**Figure 4 - heatmaps**). However, 16HBE cells displayed a higher basal SPREAD frequency compared to HBE1 cells. Against this higher baseline frequency, only TNF-α or Poly I:C induced a significant increase in SPREAD frequency, while IL-6 and IL-1β did not. (**Figure 4H**). Additionally, TNF-α or Poly I:C treatment caused SPREAD duration to increase significantly, lasting 56 minutes on average, while in all other conditions SPREADs lasted 25 to 30 minutes on average (**Figure E5A**).

**Figure 4.**
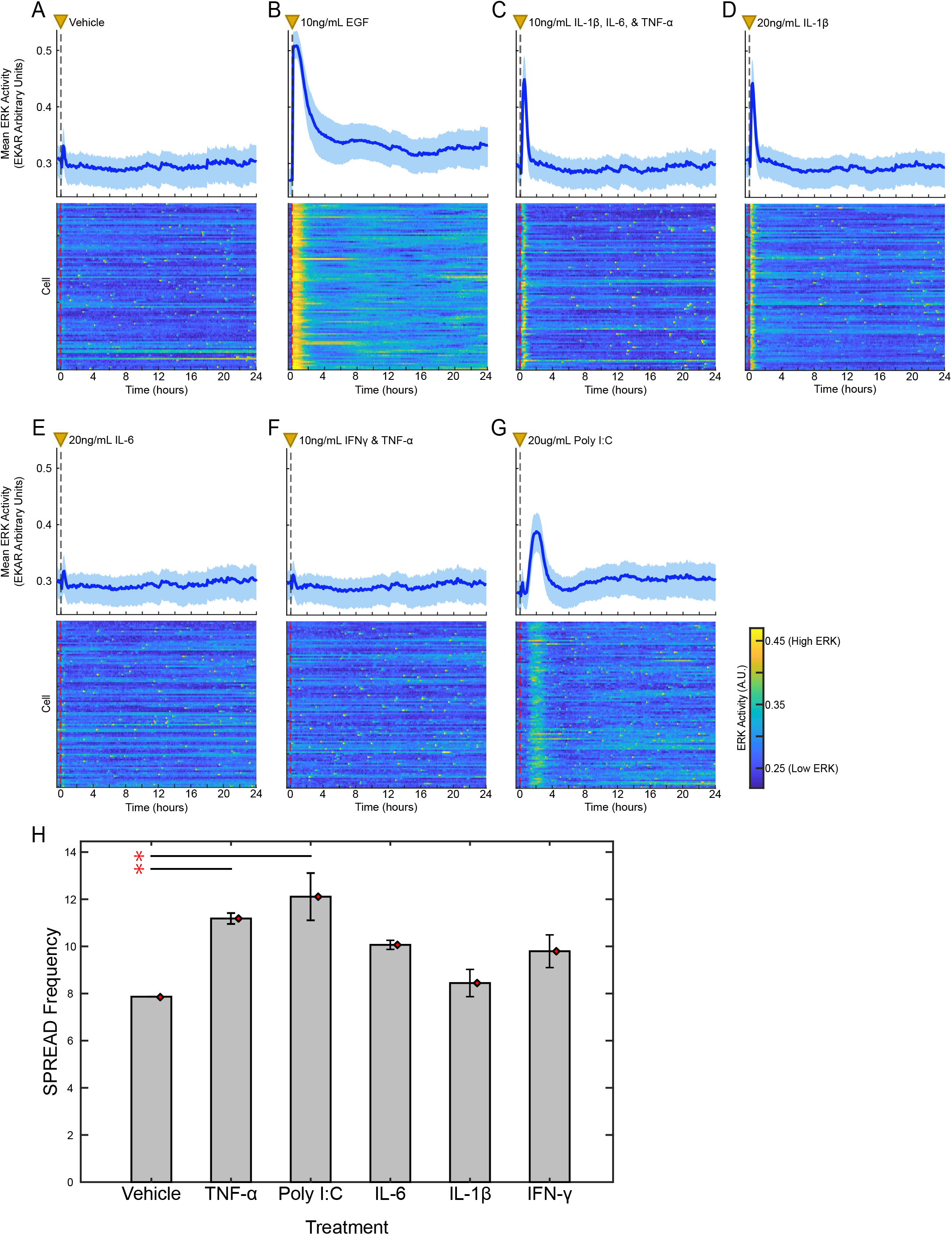
Pro-inflammatory ligands increase ERK activity in 16HBE cells. **(A-G)** Mean and interquartile range (25^th^/75^th^) of EKAREN4 signal in response to vehicle, EGF, or pro-inflammatory cytokines treatment over time. Dashed lines with yellow triangles indicate treatment (time 0 hour). **(H)** SPREAD frequency (SPREAD occurrences per hour, normalized to mm^2^) within the 24 hours following treatment. Gray bars show the mean, and red diamonds the median, of three technical replicates. Error bars show S.E.M. Red asterisks indicate conditions where SPREAD frequency is significantly different compared to the control group (P < 0.005), calculated via 1-way ANOVA compared to the control and adjusted for multiple comparisons via Dunnett’s procedure.

The difference in baseline ERK activity between HBE1 and 16HBE cells’ behavior could arise from various factors, as the two cell lines differ in their gene expression, in their ability to differentiate into ciliated epithelial cells, and in their standard growth medium composition (21, 23). Notably, HBE1 growth medium contains hydrocortisone, insulin, cholera toxin, and transferrin while 16HBE growth medium does not. These components could feasibly impact the behavior of inflammatory conditions. By visual inspection, we also noted that many of the SPREADs in 16HBE cells appeared to be initiated by apoptosis events (**Sup. Video 4**). Consistent with this observation, gefitinib treatment, which blocks apoptosis-induced SPREADs in other systems, ablated basal SPREADs observed in 16HBE cells (**Figure E5B**). Overall, results from 16HBE cells corroborate the modulation of SPREADs by inflammatory conditions as observed initially in HBE1 cells, notwithstanding several differences that are potentially explainable by the variations in culture conditions.

### Pro-inflammatory cytokines cause spatially coordinated ERK signaling in primary human airway epithelial cells

To extend our findings to a more physiologically representative setting, we introduced the ERK biosensor to primary human bronchial epithelial (pHBE) cells and measured their response to control and pro-inflammatory conditions. As expected, pHBE cells in submerged culture (i.e. not differentiated in ALI culture) did not exhibit a response to vehicle control, but displayed a strong ERK response to EGF (10 ng/mL), which validated biosensor function in primary human airway epithelial cells (**Figure 5A&B**). pHBE cells in submerged culture also displayed transient ERK activation in response to IL-1β/IL-6/TNF-α (10 ng/mL each), IL-1β (20 ng/mL), IL-6 (20 ng/mL), and Poly I:C (20 μg/mL) (**Figure 5C-G, Figure E6E**). However, pHBE cells did not respond to IL-4/5/9/13, IL-17a/23, or LPS, as seen in both immortal cell lines (**Figure E6**) We noted that IL-1β treatment caused spatially coordinated wave-like ERK activation across the entire image field, lasting 4 hours on average (**Sup. Video 5**). These large waves were not quantifiable as SPREADs since they usually do not originate within the image frame, but they suggest that primary airway epithelial cells respond in an even more coordinated manner than the cell line models.

**Figure 5.**
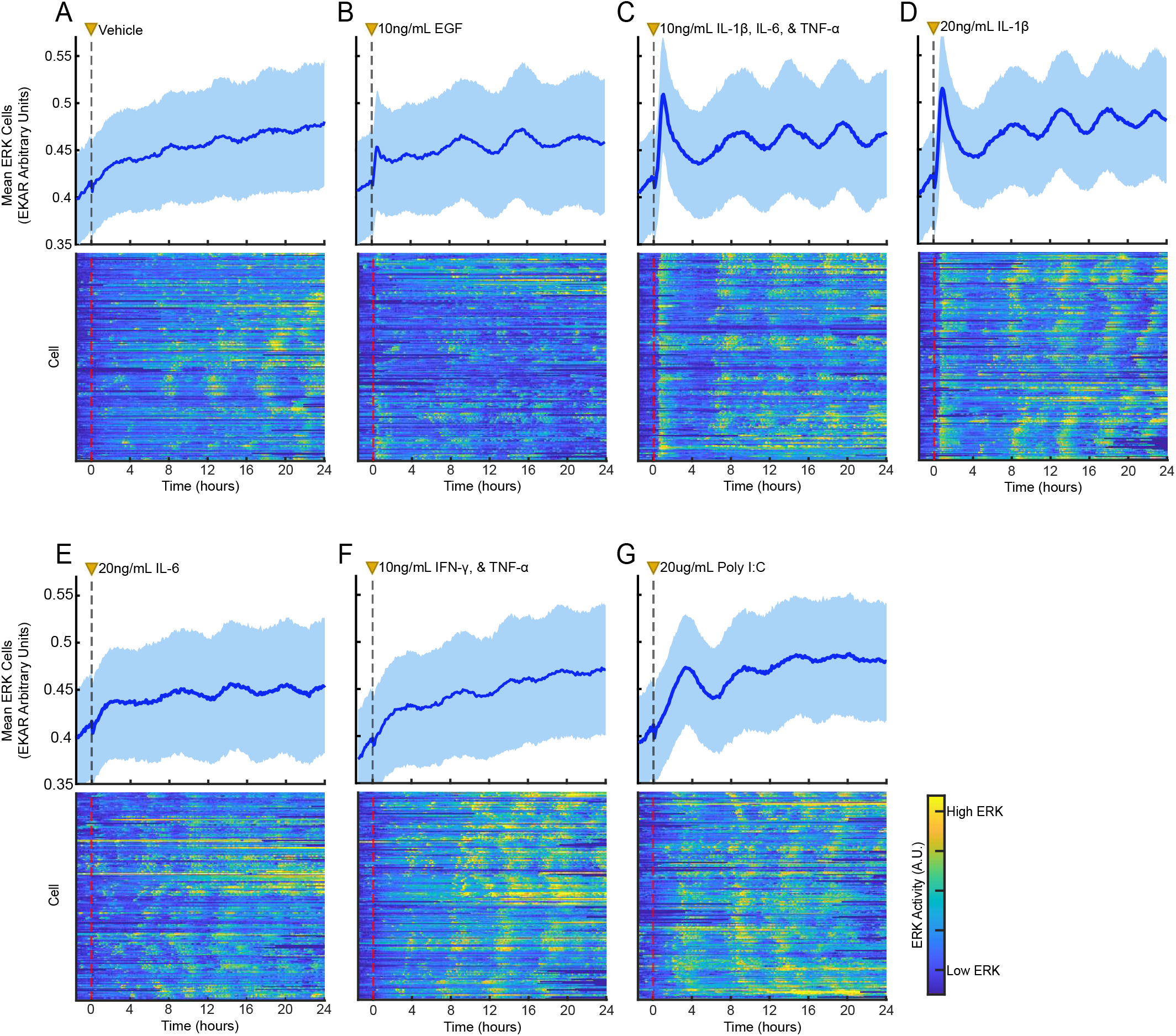
ERK responses of pHBE* cells following treatment with vehicle, EGF, or pro-inflammatory ligands. **(A-G; *top rows*)** Mean and 25^th^/75^th^ quantiles of ERK activity across all cells in given treatment from representative experimental replicates (>4,000 cells per condition across 4 technical replicates). **(A-G; *bottom rows*)** Heatmaps showing normalized ERK activity (color-axis) from 200 cells within a representative technical replicate, organized such that each cell (row) is positioned next to the closest neighboring cell in physical space. Treatment occurs at hour 0 as indicated by the dashed vertical line and yellow triangle. *primary human bronchial epithelial (pHBE) cells in submerged culture.

To investigate the effect of SPREADs on airway epithelial barrier permeability, we measured the ERK response of primary cells after differentiation in air liquid interface culture (ALI) for >30 days; differentiation was confirmed by the presence of beating cilia (**Sup. Video 6**). When treated with 20 ng/mL IL-1β, ALI-differentiated pHBE cells displayed an immediate, uniform ERK response that lasted 2 hours (**Figure 6 - Timelapse Images, Sup. Video 7**). Subsequently, SPREADs were clearly observed in all conditions, with an increased frequency of SPREADs when cells were treated with IL-1β compared to control (**Figure 6C**). Measuring the airway barrier permeability in these samples, we observed a decrease of 19% in epithelial barrier resistance as compared to control, which was not seen in EGF treated cells (**Figure 6D**). Across all samples analyzed from multiple experimental repeats, we observed a significant negative correlation between SPREAD frequency and the corresponding epithelial resistance in the same samples (R^2^=0.94, p=0.001, Figure 6E) This correlation is consistent with our hypothesis that pro-inflammatory cytokines cause increased airway barrier permeability via increased SPREAD events.

**Figure 6.**
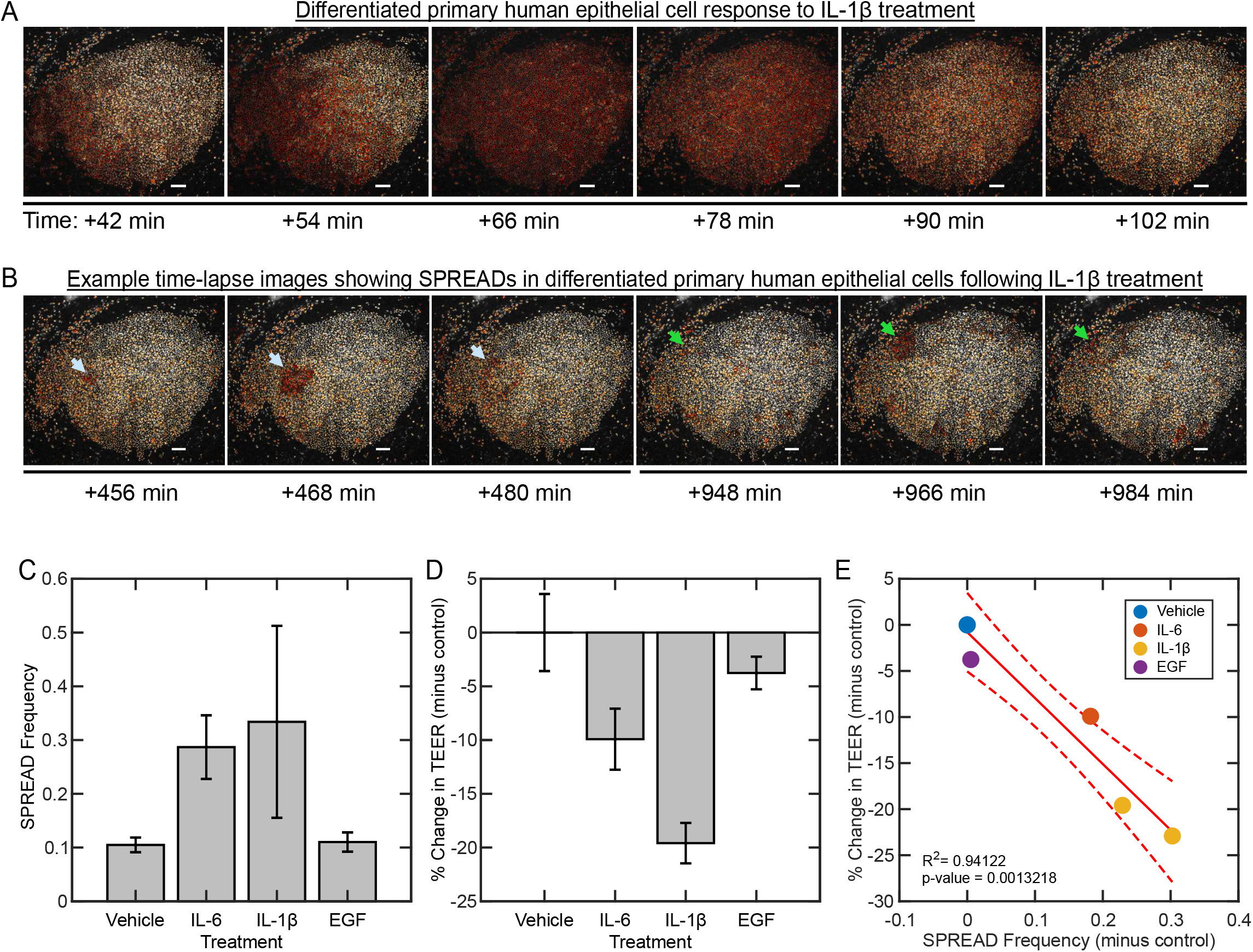
ERK responses to pro-inflammatory conditions in ALI-differentiated pHBE** cells. **(A)** Representative images showing EKAREN4 biosensor activity in response to treatment with 20 ng/mL IL-1β. **(B**) Representative image showing SPREADs occurring in differentiated pHBE cells following treatment with 20 ng/mL IL-1β. Light blue and green arrowheads identify the edge of two unique SPREAD events occurring over the time points shown. For both A and B, relative ERK activity is represented in pseudo-coloring as in **Figure 1**, with low relative ERK activity in white and high ERK activity in red. Time labels are relative to treatment with 20 ng/mL IL-1β at 0 minutes. Scale bar is 100 μm. **(C)** SPREAD frequency following treatment with indicated ligands. Red diamonds show the median of the data, and error bars show S.E.M. **(D)** Vehicle subtracted percent change in trans-epithelial resistance (TEER) following 24-hours of treatment with controls or pro-inflammatory cytokines. **(E)** Linear correlation plot of the percent change in TEER versus SPREAD frequency (both measurements are vehicle subtracted). **pHBE cells grown in bi-phasic air-liquid interface (ALI) conditions.

### Secreted IL-6 signaling plays a significant role in IL-1beta-mediated SPREADs

To establish which secreted ligands generate SPREADs, we measured cytokine abundance in conditioned media from HBE1 cells treated with IL-1β (20 ng/mL) for 12 hours. In cytokine microarray assays, GROɑ, IL-17A, IL-6, IL-8, MCP-1, IL-8, and MIP3ɑ, were all increased at least 1.5-fold by IL-1β treatment (**Figure 7A**). We cross-referenced this list of cytokines with RNA-seq data from HBE1 cells to identify ligand-receptor pairs that might cause downstream ERK activation, and thereby SPREADs, when secreted (unpublished data). Our analysis showed that IL6R and gp130 had the highest expression amongst the receptor candidates, prioritizing IL-6 as a potential candidate that is further supported by multiple reports in the literature (31, 32). Bead-based cytokine assays further confirmed IL-6 secretion in IL-1β-treated HBE1 cells (**Figure E7**).

**Figure 7.**
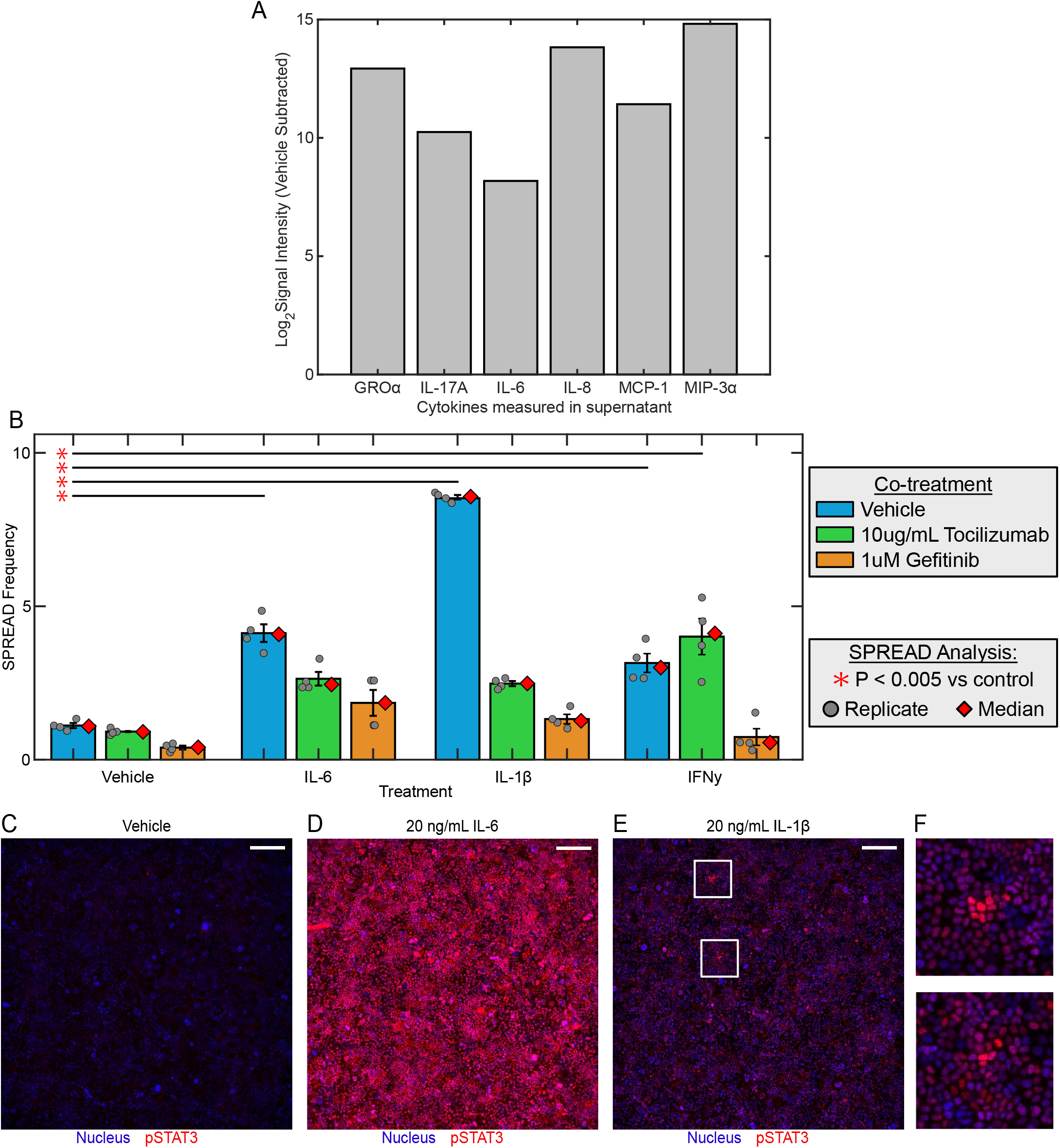
Investigating the mechanism of cytokine-mediated SPREADs. **(A)** Vehicle-subtracted Log2 mean intensity of secreted cytokines in the supernatant of IL-1β treated HBE1 cells versus control. **(B)** SPREAD frequency of HBE1 cells co-treated with vehicle or pro-inflammatory conditions (indicated by labels under bars) plus vehicle, tocilizumab, or gefitinib (indicated by bar color). **(C-E)** pSTAT3 immunofluorescence (IF) stains of HBE1 cells after 24-hours treatment with vehicle, 20 ng/mL IL-6, or 20 ng/mL IL-1β. The nuclear marker is shown in blue with pSTAT3 shown in red. All image intensities are scaled similarly per channel. White boxes outline regions of high pSTAT3 activity in IL-1β treated cells. Representative images selected from 9 technical replicates across 3 independent experiments. Scale bars in panels C-E are 250 μm. **(F)** Enlarged images of clusters of pSTAT3 activation, identified in panel E by two white boxes. For panel F, the whole image in panel E was scaled 4x then cropped.

As IL-6 can activate ERK through IL6R mechanisms that are EGFR-dependent (33, 34) and EGFR-independent, (33, 35) we sought to determine if the mechanism of ERK activation in SPREADs was unimodal (acting through EGFR or IL6R alone) or multimodal (via multiple receptors). We treated cytokine-exposed HBE1 cells with inhibitors of EGFR or IL6R (gefitinib and tocilizumab, respectively; **Figure 7B**). HBE1 cells treated with IL-1β and gefitinib or tocilizumab displayed significantly fewer SPREAD events, compared to IL-1β alone. This effect suggested that IL1β-mediated SPREADs are caused via both EGFR and IL6R ligands. However, IFN-γ-induced SPREAD frequency was reduced significantly by gefitinib, but not by tocilizumab, indicating that IFN-γ-mediated SPREADs are primarily EGFR-dependent (**Figure 7B**). Interestingly, tocilizumab or gefitinib treatment significantly decreased IL-6-induced SPREADs, further implying that some SPREADs depend on both EGFR and IL6R activity (**Figure 7B**).

To further investigate the role of IL-6 in spatially localized signaling events, we performed immunofluorescence for phosphorylated STAT3 (pSTAT3), the transcription factor canonically activated by IL-6. As expected, we observed minimal pSTAT3 in control-treated HBE1 cells (**Figure 7C**) while IL-6-treated conditions displayed uniform high STAT3 activation (**Figure 7D**). However, under SPREAD-promoting conditions (IL-1β or IFN-γ), we observed clusters of cells with STAT3 activation which were similar in size and shape to SPREADs (**Figure 7E&F; Outlined by white boxes**). Together, the secretion of IL-6, the dependence of SPREADs on IL6R, and the co-occurence of pSTAT3 indicate that IL-1β-induced SPREADs in airway epithelial cells are markedly dependent on IL-6 signaling, in addition to the known role of EGFR found in other epithelia.

### Metabolic suppression limits cytokine-induced SPREAD frequency

Patients with inflammation-related metabolic syndromes, including diabetes and obesity, show reduced capacity for mucosal barrier repair and are at greater risk for airway barrier-related inflammation (36–39). Considering this comorbidity, we investigated the impact of metabolic regulators on cytokine-induced SPREADs in our HBE1 cells. To verify elevated metabolic stress, we introduced a second fluorescent biosensor for AMP-activated protein kinase (AMPK) into our HBE1 cells expressing the ERK biosensor (**Figure 8A**). AMPK responds to changes in cellular energetic status and may limit cytokine production when active (40), potentially suppressing cytokine-induced SPREADs.

**Figure 8.**
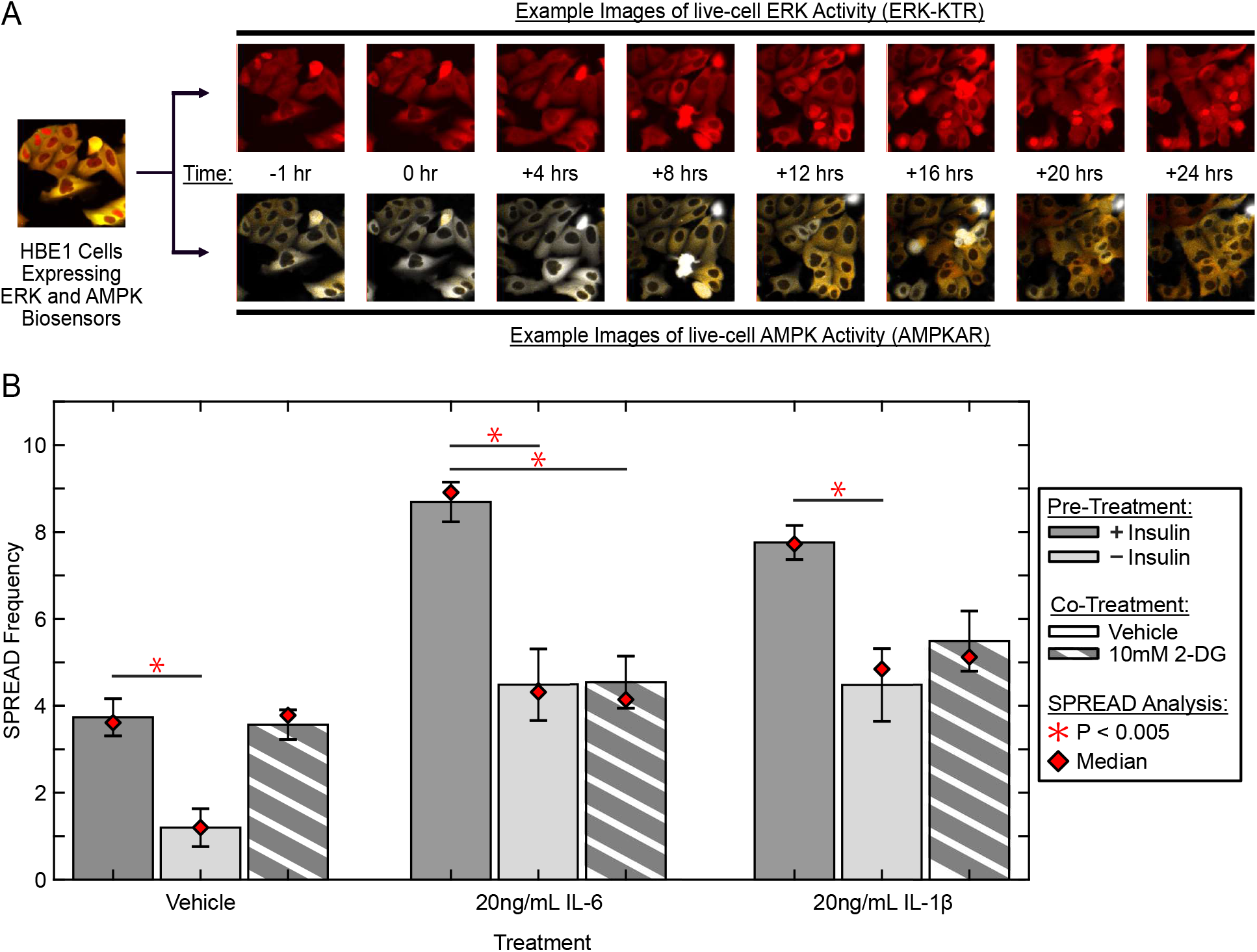
Metabolic regulation of cytokine-mediated SPREADs. **(A)** Example images of HBE1 cells expressing both the ERK and AMPK biosensor. *Top* images show ERK activity over time and bottom images show AMPK activity over time in the same cells. Cells in example images were treated with EGF and 2-deoxyglucose (2-DG) at 0 hours. **(B)** SPREAD frequency of HBE1 cells treated with pro-inflammatory cytokines following 16 hours of either starvation or supplementation of insulin, or co-treatment with either vehicle or 10mM 2-DG. Red diamond is the median of data, error bars are S.E.M. Data are 4 technical replicates across 2 experimental replicates.

We first explored how insulin deprivation affected cytokine-induced SPREADs and AMPK activity in our HBE1 cells. Insulin deprivation significantly reduced the frequency of pro-inflammatory cytokine-induced SPREADs (**Figure 8B**). Specifically, we noted that insulin-starved cells treated with IL-1β (10 ng/mL or 20 ng/mL), or IL-6 (20 ng/mL) displayed 25%, 42%, and 48% fewer SPREADs relative to their insulin-treated counterparts. This suppression decreased SPREAD frequency to no longer be significantly different than vehicle. To further investigate the idea that metabolic stress decreases cytokine-induced SPREADs, we treated the cells with 2-deoxyglucose (2-DG), an inhibitor of glycolysis, to cause maximal metabolic stress. 2-DG treatment induced maximal AMPK activity, indicating high metabolic stress in the cells (**Figure E8**). 2-DG treatment led to a significant decrease in IL-6-, high IL-1β-, and low IL-1β-induced SPREAD frequency, when compared to the vehicle-treated counterparts for each cytokine (**Figure 8C**), further indicating that increased metabolic activity correlates with higher SPREAD frequency.

### Corticosteroids suppress SPREADs

Corticosteroids such as hydrocortisone are common anti-inflammatory drugs used clinically (41, 42). Their activity is attributed to multiple mechanisms, including suppressed secretion of IL-6 and other pro-inflammatory cytokines (43, 44) and reduced glycolytic metabolism (45). As our previous results implicate both IL-6 secretion and glycolytic metabolism in the promotion of SPREADs, we tested the effect of HC on cytokine-induced SPREADs. We compared HBE1 cells pretreated for 24 hours with 0.5 μg/mL HC (1.38 μM) to cells grown without HC for 24 hours. HC-treated cells were much less susceptible than untreated cells to SPREADs induced by TNF-α and IL-1β with reductions in SPREAD frequency of 80% and 71%, respectively (**Figure 9A**). Together with the higher SPREAD frequency found in 16HBE cells, which as noted earlier are cultured in the absence of HC, these data indicate that HC potently suppresses cytokine-mediated SPREAD activity.

**Figure 9.**
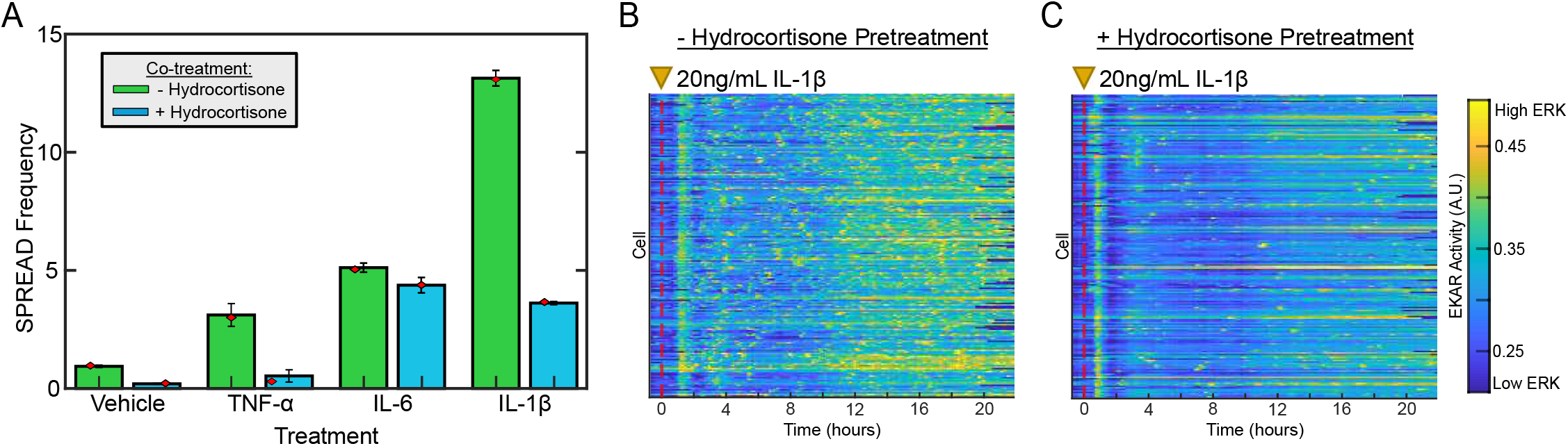
Corticosteroids suppress pro-inflammatory cytokine-induced SPREADs in HBE1 cells. **(A)** Suppression of TNF-α, IL-6, and IL-1β -induced SPREADs by 24-hour pre-treatment with 1.38 μM hydrocortisone and cytokine treatment at 20 ng/mL, each) **(B and C)** Heatmaps showing ERK activity (color-axis) from 200 cells within a representative technical replicate organized such that each cell (row) is positioned next to the closest neighboring cell in physical space. 20 ng/mL IL-1β treatment occurs at hour 0 as indicated by the yellow triangle and dashed line.

## DISCUSSION

There is a pressing need to better understand inflammatory mechanisms in airway epithelial cells that lead to chronic disease pathogenesis and progression. Our data offer deeper insight into the long-standing question of ERK’s dual role as a pro- and anti-inflammatory signaling kinase, revealing for the first time intricate spatial patterns of ERK activity (SPREADs) within airway epithelial cells. Specifically, we establish that pro-inflammatory environments promote spatially *localized* ERK activity in the form of SPREADs, whereas growth factor treatment causes more uniformly *distributed* ERK activity across all measured airway epithelial cells. This difference indicates a spatiotemporal specialization of ERK activity that is in part dependent on the stimulatory context.

We identify pro-inflammatory conditions including IL-1β, IL-6, TNF-α, and IFN-γ, that increase the frequency of SPREADs through secreted EGFR and IL6R ligands. Our findings suggest a novel mechanism by which cytokine-mediated ERK activity can cause collective cell behavior in the airway epithelium that differs from canonical EGF-mediated ERK signaling. We propose that localized SPREAD events in specific areas caused by exposure to pro-inflammatory ligands may be a mechanism for targeted ERK-mediated airway barrier permeabilization. SPREADs caused by localized cytokine secretion in the lung epithelium could synchronize barrier permeability (5, 12) with cytokine-mediated chemoattraction of innate and adaptive immune cells at specific sites, providing an entry mechanism for immune cells (**Figure 10**). This proposed mechanism is distinct compared to the barrier-protective effects of EGF-mediated ERK activity (46), which we demonstrate to be uniformly distributed in epithelial monolayers. Localized changes in epithelial plasticity mediated by ERK have also been observed in Drosophila and zebrafish models (47, 48).

**Figure 10.**
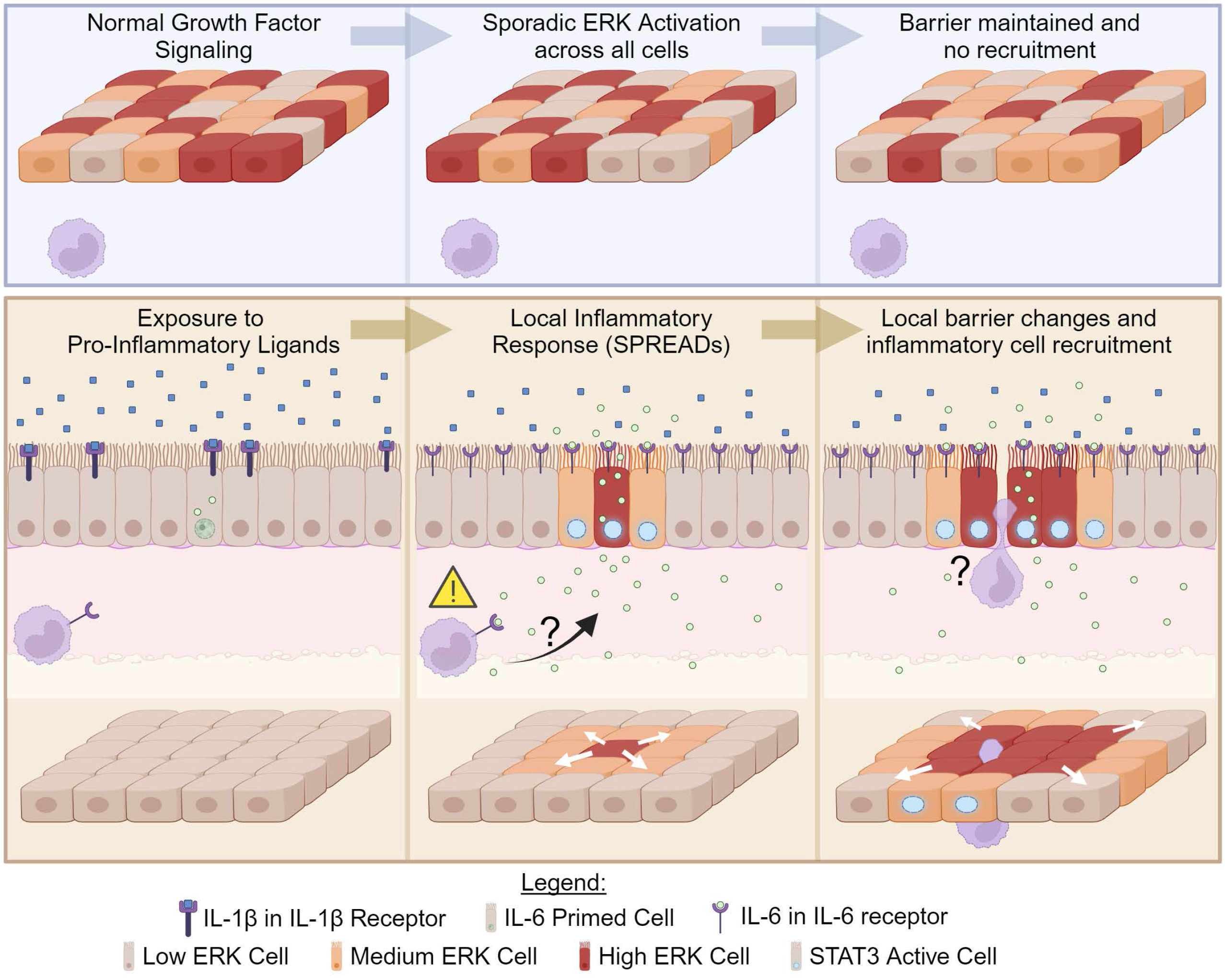
Model for the effects of pro-inflammatory cytokines on SPREADs in the airway epithelium. *(Top row - light blue box)* Growth factors such as EGF induce sporadic ERK activation uniformly across the epithelium. Without strong spatial organization, this signaling does not induce rapid morphological changes in the cell layer or create a focal source of cytokine secretion that would recruit innate immune cells. (Bottom row - beige box) Exposure to pro-inflammatory ligands such as IL-1β triggers secretion of ligands such as IL-6 in a small number of cells, initiating SPREADS. We hypothesize that this organized wave-like ERK and STAT3 activity drives local changes in the structure of the epithelium, promoting small zones of permeability. We further hypothesize that the gradient of IL-6 (and other cytokines) generated by this focal secretion recruits innate immune cells such as neutrophils (purple) while simultaneously inducing permeability to permit their passage through the epithelium. Such a model could explain the contrasting and context-dependent roles that have been proposed for ERK in the airway epithelium, either (i) as a promoter of epithelial integrity under uniform stimulation or (ii) as a promoter of inflammation under the newly described SPREAD-like stimulation. Figure created with BioRender.com.

Our study adds to the growing body of literature that underscores the importance of spatial signaling in maintaining proper epithelial homeostasis. A low frequency of SPREADs is important for epithelial homeostasis, specifically tuning cell survival and apoptosis rates to the local context of each cell and allowing the tissue to adapt to cell death (19) and division events (18). On the other hand, continuous high frequency SPREAD events, such as those caused by pro-inflammatory cytokines, might have deleterious effects on the epithelium by triggering a positive feedback loop of epithelial permeability and immune cell recruitment. Beyond ERK, multiple pathways could exhibit SPREAD-like behavior and collectively influence tissue systems. For example, calcium waves from Caspase-8-expressing cells or a laser-ablated apoptotic cell in zebrafish promote RAS-mutant MDCK cell ejection (49). Locally released cytokines such as IL-1β or TNF-α can propagate signals between immune cells and epithelial cells through the NF-κB and JNK pathways (50). Collectively, these ligands, receptors, and pathways allow for a complex spatiotemporal flow of information regulating cell behavior. Our study highlights how methods with high spatial and temporal resolution can be used to understand how these pathways operate in multicellular systems such as the airway epithelium.

Our observations that cytokine signaling, hydrocortisone, and metabolic conditions all modulate SPREAD frequency are consistent with their known roles in inflammation. However, we also provide for the first time evidence that these factors operate, at least in part, through the shared mechanism of ERK SPREADs. This convergence suggests that future studies evaluating possible synergistic effects of anti-cytokine drugs, metabolic inhibitors (e.g., metformin), and anti-inflammatory drugs (e.g., corticosteroids, statins) may reveal combinations of treatments with clinical utility in a manner not previously imagined. It may also be possible to identify clinical alternatives for patients who are resistant to SPREAD modulation through one mechanism (e.g., corticosteroids) but sensitive through another (e.g., EGFR/IL6R inhibitors, metformin, statins, phosphodiesterase-4 inhibitors, leukotriene receptor antagonists, etc.). While biosensors cannot be used directly in human patients, they could feasibly be applied to primary human airway epithelial tissue from donors. SPREAD mechanisms could thus be assessed in individual patients as part of a precision medicine approach to the mucosal innate immune response.

In conclusion, our findings indicate that pro-inflammatory cytokines induce SPREADs in airway epithelial cells, and that SPREADs play an important role in coordinating the response of airway epithelial cells to changes in their environment, particularly in conditions where airway epithelial barrier breakdown is observed. These findings contribute to our understanding of the complex signaling networks that regulate the response of lung epithelial cells during inflammation, and show a novel mechanism of airway disease progression, and provide insights into potential therapeutic targets for airway diseases associated with ERK signaling and barrier dysfunction.

## ACKNOWLEDGEMENTS

NLD wishes to thank the UC Davis Heath Comparative Lung Biology and Medicine Training Grant (T32) and its members for training on writing and general feedback on the manuscript. Cell sorting for this project was provided by the University of California Davis Flow Cytometry Shared Resource Laboratory which is funded by the NCI P30 CA093373 (Comprehensive Cancer Center), and S10 OD018223 (Beckman Coulter “Astrios” cell sorter and “Cytoflex” cytometer) grants, with technical assistance from Bridget McLaughlin and Jonathan Van Dyke. We also wish to thank Abhineet Ram for sharing the base MATLAB code adapted for making spatial heatmaps.

## Author Contributions

NLD - First to discover and discuss SPREADs in HBE cells in response to cytokines, helped conceive project, designed and ran experiments, performed image analysis on experiments, helped validate and improve our matlab implementation of ARCOS, did statistical analysis on data, made figures, and wrote manuscript.

DPO - Helped make and our matlab implementation of ARCOS, supported updates and upgrades to our matlab implementation of ARCOS, helped write methods section, edited paper, checked and edited figures.

KJC - Provided HBE1 and pHBE cells, helped with experimental optimization and design, set up and cultured pHBE cells in ALI with EKAREN4 biosensor, helped run TEER measurements, performed cytokine luminex assay, edited manuscript.

MP - Helped with image analysis, ERK pulse detection, build and validate our matlab implementation of ARCOS, determine statistical analysis used, and validate findings. Contributed to manuscript writing, editing, and review.

JMF - Made HBE1 and 16HBE EKAREN4 cell lines, set up experiments and cell culture, helped collect and process data, and reviewed the manuscript text.

DM - Conducted pSTAT3 immunofluorescence staining, performed and measured the ELISA cytokine assay.

AAZ – Co-conceived of the project and basic idea to assess spatio-temporal signaling in submerged epithelial cells and fully differentiated human airway epithelial cells, helped design experiments, analyze and interpret data, write and edit the manuscript.

JGA - Co-conceived of the project and basic idea to assess spatio-temporal signaling in submerged epithelial cells and fully differentiated human airway epithelial cells, helped design experiments, analyze and interpret data, write and edit the manuscript.

## Funding Sources

<u>American Lung Association:</u>

IA-628171 (JGA)

<u>National Heart, Lung, and Blood Institute (NHLBI):</u>

HL007013 (NLD) (UC Davis Lung Center T32)

R01 HL148715-01A1 (AAZ)

R01 HL151983-01A1 (JGA)

## Online Data Supplement

### SUPPLEMENTAL METHODS

#### Cell culture and Media

##### HBE1 Cells

HBE1, a papilloma virus-immortalized human bronchial epithelial cell line originally generated by Dr. James Yankaskas (University of North Carolina, Chapel Hill, NC) (E1), was provided to our lab via Dr. Reen Wu of University of California, Davis. Routine cell culture for the HBE1 cell line was performed as described (E2) and maintained in ‘HBE1 Culture Medium’ with the modification of Hydrocortisone used in place of Dexamethasone (see table E1 - HBE1 Culture Medium). All cells were routinely passaged prior to reaching confluence and were between passage 8 and 28.

##### 16HBE Cells

16HBE14o-(referred to as 16HBE) cells were obtained from Sigma-Aldrich (#SCC150) and cultured in ɑ-MEM (Sigma M2279) + 2mM L-Glutamine + 10% FBS (E3). Again, all cells were routinely passaged prior to reaching confluence and were between passage 3 and 15.

##### Primary Human Bronchial Epithelial (pHBE) Cells

Primary Human airway epithelia (pHBE) cells were used for submerged experiments immediately after transduction with EKAREN4 or cultured for at least 4 weeks in an air-liquid interface culture system as described by Wu and colleagues before use in live-cell experiments (E4). Briefly, cryopreserved pHBEs were centrifuged at 280 × g for 5 min and resuspended to 5×10^4^ cells/mL in complete Bronchial Epithelial Cell Growth Medium (BEGM; Lonza Group Ltd. CC-3171); were seeded to 100 mm dishes. Upon initial plating of 500,000 cells per 100 mm dish cells were maintained in pHBE cell expansion medium for 5 - 6 days of proliferation (See table E2 Primary Human Airway Culture Cell Expansion Medium). At near confluency, cells were lentiviral transduced with the EKAREN4 biosensor (ERK biosensor) and subsequently used for live-cell experiments or passaged to air-liquid-interface inserts for long-term culture and live-cell imaging after differentiation occurred (>30 days). Near-confluent cells were sub-cultured into air-liquid interface (ALI) culture inserts (Corning 3460, 12 well) at a ratio of 30 wells per 100 mM dish and maintained in primary human airway culture differentiation media (See table E3; without all-trans-retinoic acid). Apical media was removed after two days of confluency and basal well media was supplemented with all-trans-retinoic acid (30 nM, Sigma R2625) for continuation of culture. Confluence for over 10 days required media changes twice per day, due to nutrient depletion, and maintained up to 10 weeks for high-density culture (2 x 10^6^ cells per cm^2^) with copious beating cilia and flowing mucous channels.

#### Biosensor Delivery and Usage

ERK biosensors were stability integrated into HBE1, 16HBE, and pHBE cells via the lentiviral expression system using the pLJM1 vector. Example images of ERK biosensors (ERKTR (E5) and EKAR (EKAREN4) (E6)) in HBE1 cells are shown in **Figure 1A**. Biosensor designs were previously described in (E7) (referred to as: ERK-KTR-IRES-NLS-mCardinal-Hygro (ERKTR) and EKAR-EN4-NLS-Puro (EKAREN4)). AMPK biosensor was stably integrated into HBE1 cells already expressing ERK-KTR via piggyback transposase mediated integration and were delivered via NEON electroporation following manufacturer instructions. After transduction, immortal cell lines (HBE1 and 16HBE) expressing biosensors were isolated in bulk or clonally using flow cytometry sorting. For each biosensor, we isolated stable cells with homogenous expression; all data reported in this study reflect representative behaviors that were consistent across cell line populations and passages used. All cell lines were confirmed to be mycoplasma-negative by third-party testing (ATCC). Primary cells were not sorted to avoid unnecessary cell stress.

#### Live-cell Fluorescence Microscopy

Time-lapse wide-field microscopy was performed as described previously, with the modification of initial cell seeding and days seeded prior to use (E8, E9). Briefly, 20,000-30,000 cells (667-1000 cells / mm^2^ culture surface area) were seeded into each well of a glass-bottom 96-well plates (Cellvis P96-1.5H-N, Mountain View, CA) that were treated with type I collagen (GIBCO A10483-01) to promote cell adherence, two days prior to imaging. Cells were visually verified to be at 100% confluence via brightfield microscope prior to imaging.

For live-cell microscopy experiments, cells were washed two times with and then cultured in their respective ‘Imaging Medium’ for at least 6 (up to 24) hours prior to imaging. ‘Imaging Medium’ is a custom media formulation, similar to GIBCO 11320-033 DMEM/F12 or Sigma M2279 - ɑ-MEM, however it is formulated to be lacking glucose, glutamine, pyruvate, riboflavin, folic acid, and phenol red (UC Davis Veterinary Medicine Biological Media Service), which could be supplemented for experimental use. Before imaging, HBE1 cells with the ERK-KTR/AMPKAR2 biosensors were treated with 0.1 ug/mL Hoechst-33342 (Life Technologies: H3570) to maximize nuclear identification and cell tracking. In later experiments with HBE1, 16HBE, and pHBE expressing the nuclear localized EKAREN4 biosensor, cells were not treated with Hoechst, as it is unnecessary.

During experiments, cells were maintained in an environmental chamber in 95% air and 5% CO2 at 37 C. Fluorescent images were collected with either a Nikon (Tokyo, Japan) 20X/0.75 NA Plan Apo objective or 10X/0.4 NA Plan Apo objective on a Nikon Eclipse Ti2 inverted microscope, equipped with a Lumencor SPECTRA III or Lumencor SPECTRA X light engine. Fluorescence filters used in the experiment are: BFP/DAPI (custom ET395/25x - ET460/50 m - T425lpxr, Chroma), CFP (49001, Chroma), YFP (49003, Chroma), Cherry (custom FF01-586/20 – FF01-640/40 – FF605-Di02, Semrock) and Cy5 (49006, Chroma). The DAPI filter was used for Hoechst-33342 containing experiments, CFP and YFP filters were used for AMPKAR2 or EKAREN4 experiments, the Cherry filter was used for ERK-KTR experiments images, and the Cy5 filter was used for pSTAT3 stain images. Images were acquired using Andor Zyla 5.5 sCMOS or Teledyne Photometrix Kinetix sCMOS camera every 3 or 6 minutes. Exposure times for each channel were 50-75 ms for DAPI; 75 – 300 ms for CFP; 150 – 350 ms for YFP; 150 – 400 ms for Cherry and 50-250 ms for Cy5 and ranged in power from 5-50% for the channels depending on reporter expression and light source.

All experiments involving HBE1 cell lines were performed in ‘HBE1 imaging medium’, unless otherwise indicated (see media composition table E4 - HBE1 Imaging Medium). HBE1 imaging medium is similar to normal HBE1 cell culture medium, however it is lacking EGF and BNE. HBE1 imaging medium was supplemented with glucose, glutamine, pyruvate, insulin, transferrin, cholera toxin, and hydrocortisone, unless otherwise indicated. For HBE1 experiments testing insulin deprivation and hydrocortisone presence, the respective component was removed from the HBE1 imaging medium, with no other modification.

16HBE imaging experiments were performed in ‘16HBE imaging medium’ (see media table E5 - 16HBE imaging medium). 16HBE imaging medium is a custom media formulation, similar to Sigma M2279 - ɑ-MEM, but lacked glucose, glutamine, pyruvate, riboflavin, folic acid, and phenol red (UC Davis Veterinary Medicine Biological Media Service). 16HBE imaging medium was supplemented with glucose and L-glutamine to recapitulate their normal growth medium without FBS supplementation.

Submerged and ALI differentiated pHBE imaging experiments were performed in submerged pHBE imaging medium (see table E6) or Air-Liquid Interface pHBE imaging medium (see table E7), respectively. These imaging media are similar to their typical growth medium (see tables E2 and E3), lacking epidermal growth factor, insulin, transferrin, dexamethasone/hydrocortisone, cholera toxin, bovine hypothalamus extract, and all trans retinoic acid.

#### Cell identification, tracking, and biosensor signal quantification

Image data were processed to segment each cell’s nucleus and cytoplasm, and mean pixel intensity values were determined using a custom procedure written in MATLAB (E10). Image data are read from NIS Elements generated ND2 files using a minimally modified version of the bioformats MATLAB toolbox provided by Open Microscopy (https://www.openmicroscopy.org/) (E11).

Cell nuclei are identified and masked by binarizing images at multiple intensity thresholds, and logging approximately circular objects within expected size constraints. A donut-shaped region of each cell’s cytoplasm is masked by taking the identified nucleus and expanding that mask outwards with a gap. The mean intensity of a cell’s nucleus and its respective cytoplasm are recorded for each image channel collected. Cells are tracked over time using a MATLAB implication of uTrack 2.0 (E12). Single-cell ERK activity was determined from the ERK-KTR data by taking the ratio of the cell’s mean cytoplasmic cherry signal intensity divided by the cell’s mean nuclear cherry signal intensity. From EKAREN4, activity was calculated as 1 - (Nuclear CFP intensity / Nuclear YFP intensity) * PowerRatio; this yields a linear relationship with the fraction of reporters activated. AMPK activity was similarly calculated as 1-(Cytoplasmic CFP intensity / Cytoplasmic YFP intensity) * PowerRatio. In both cases PowerRatio is the ratio of total power collected in CFP over that of YFP (each computed as the spectral products of the relative excitation intensity, exposure time, molar extinction coefficient, quantum yield, light source spectrum, filter transmissivities, and fluorophore absorption and emission spectra of each channel). Single-cell ERK and AMPK activity was quantified, then filtered to omit data tracks that did not last at least 8 hours or showed signal intensities outside of reasonable expected ranges (KTR = 0-3.5; EKAREN4/AMPKAR2 = 0.2-0.9). Pulse annotation and quantification were performed using MATLAB, which is described in (E8, E13). Analysis of “responder” cells were defined as cells that increased in ERK-KTR or EKAREN4 signal by 0.3, or 0.05 A.U., respectively, when mean activity 1 hour before treatment was subtracted from mean signal 30 minutes after treatment.

#### Quantification of spatially localized ERK activity (SPREADs)

To quantify spatially localized ERK activity (SPREADs), we developed a MATLAB implementation of Automated Recognition of Collective Signaling (ARCOS) developed by the Pertz Lab (E14) with improvements to the binarization and clustering methodology. Following the ARCOS methodology, time series data are binarized and subsequently clustered with Density Based Spatial Clustering of Applications with Noise (DBSCAN). Cluster labels are tracked using the k-nearest neighbors algorithm to match neighbors of clusters within epsilon distance between time points.

Our implementation expanded on ARCOS by adding automatic estimations of the “epsilon” and “minpts” hyperparameters for DBSCAN, based on the distributions of the input data. Furthermore, we allow any alternative binarization method to be provided, and in this study binarize cells using “pulses” or peaks identified in ERK activity across time via the pulse analysis methodology described previously (E8). This allows for cells with heterogeneous baseline signal activity and decreases instances of false binarization seen in simpler methods that do not consider activity over time. This implementation includes automated correction of erroneously merged SPREADs (e.g. when a spread ends within one timepoint of another beginning nearby).

Cluster data can be filtered by duration, start time, maximum area, maximum count, mean rate of change of count and mean rate of change of area, effectively narrowing down the pool of spread candidates. In our case, we filtered all data to eliminate falsely identified clusters that are not SPREADs. This included eliminating SPREADs that did not include at least 3 cells, lasted under 6 minutes, or lasted for more than 3 hours.

We validated our detection approach by generating synthetic spreads and using Adjusted Rand Indices (ARI) to compare per-timepoint ground truth cluster assignments to ARCOS cluster assignments of varying lifetime and frequency. We observed mean per-timepoint ARI values between 0.96 and 0.99, indicating that our methodology is sufficiently robust to capture SPREADs with high fidelity.

#### Code and data availability

Raw image data are available on request, as they are too large to upload to a remote server with reasonable expectations for download ability.

Image recognition software used by the lab is described in (E8, E10) and is available upon request.

All code such as the scripts used to produce the figures in this paper, including raw analysis outputs and statistics along with methods are available at the following github repositories (to be made public after submission acceptance):

Our matlab implementation of arcos: https://github.com/Albeck-Lab/ARCOS-MATLAB

ERK activity and SPREAD data analysis: https://github.com/Albeck-Lab/SPREADs

An html version of all code used to make figures and their respective outputs (including statistics) is available here: https://github.com/Albeck-Lab/SPREADs/tree/main/Plotting_Code/html

## SUPPLEMENTAL FIGURE LEGENDS

**Figure E1.**
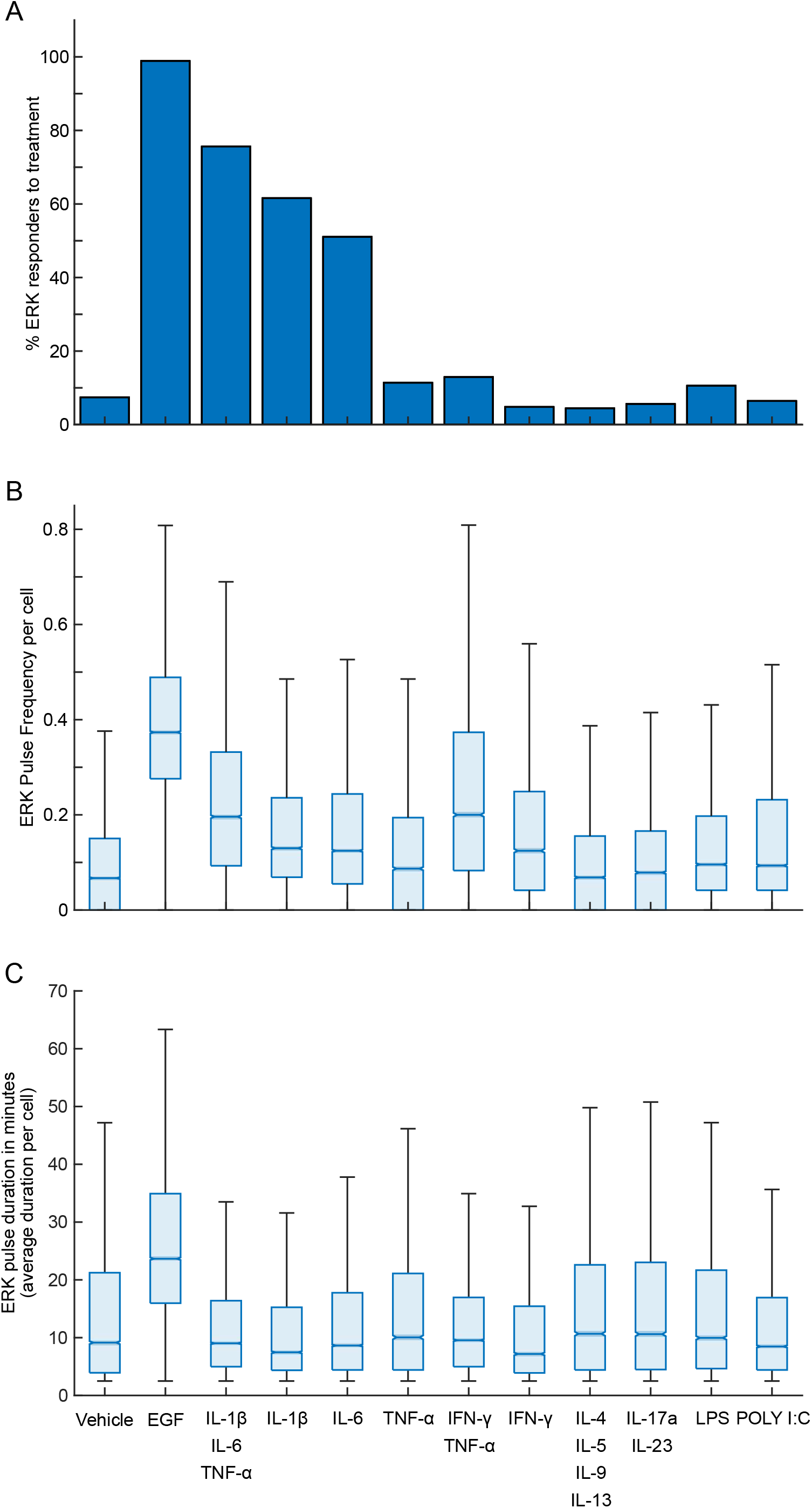
Quantification of ERK activity in HBE1 cells in response to pro-inflammatory ligands and controls. **(A)** The percentage of cells that respond to respective treatments. Responders were defined as cells that increase in ERK-KTR or EKAREN4 signal by 0.3, or 0.05 A.U., respectively, after treatment, relative to mean activity 1 hour before treatment. **(B)** Notched box plots of ERK pulse frequency over 24 hours following treatment (occurrences of ERK activation per cell per hour) for each condition. Plots represent the median (line at center of notch), 95% confidence interval of the mean (notch), interquartile range (25^th^/75^th^ percentile, blue box), and range of data excluding outliers (whiskers). **(C)** Notched box plots showing each cell’s average ERK pulse duration (in minutes). Data for A-C is cumulative from 5 experimental replicates, with >7,500 cells per condition.

**Figure E2:**
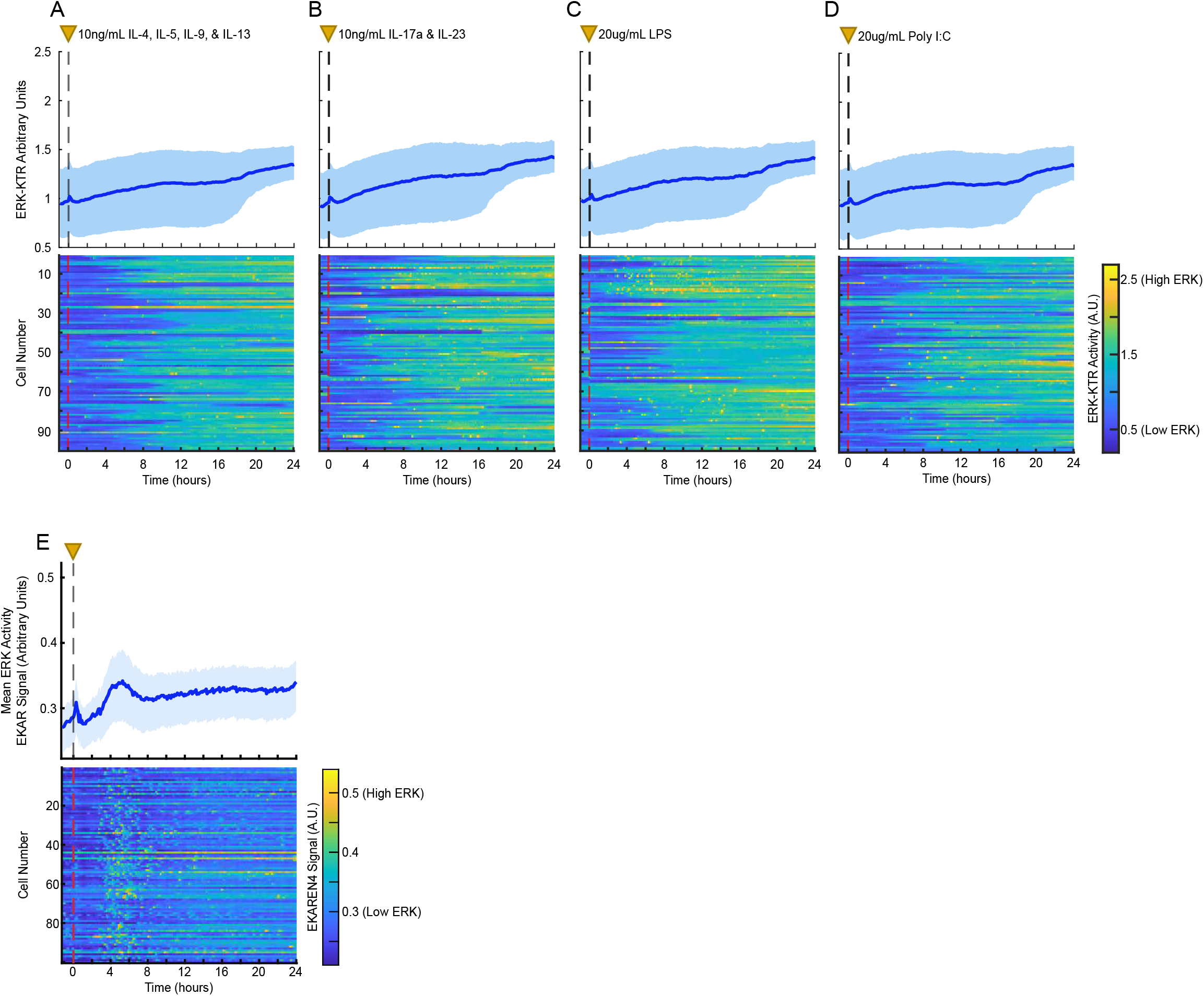
ERK responses of HBE1 cells to pro-inflammatory ligands. **(A-D)** Mean and heatmap plots showing HBE1 cell ERK activity in response to pro-inflammatory ligands and vehicle control. Treatments occur at 0 hours, indicated by a vertical dashed line and yellow triangle. Heatmaps show ERK activity (color-axis) of 100 randomly selected cells from the data that make up the mean data plots shown above. Heatmap plots are representative data from 1 of 3 technical replicates, with >500 cells per condition.

**Figure E3:**
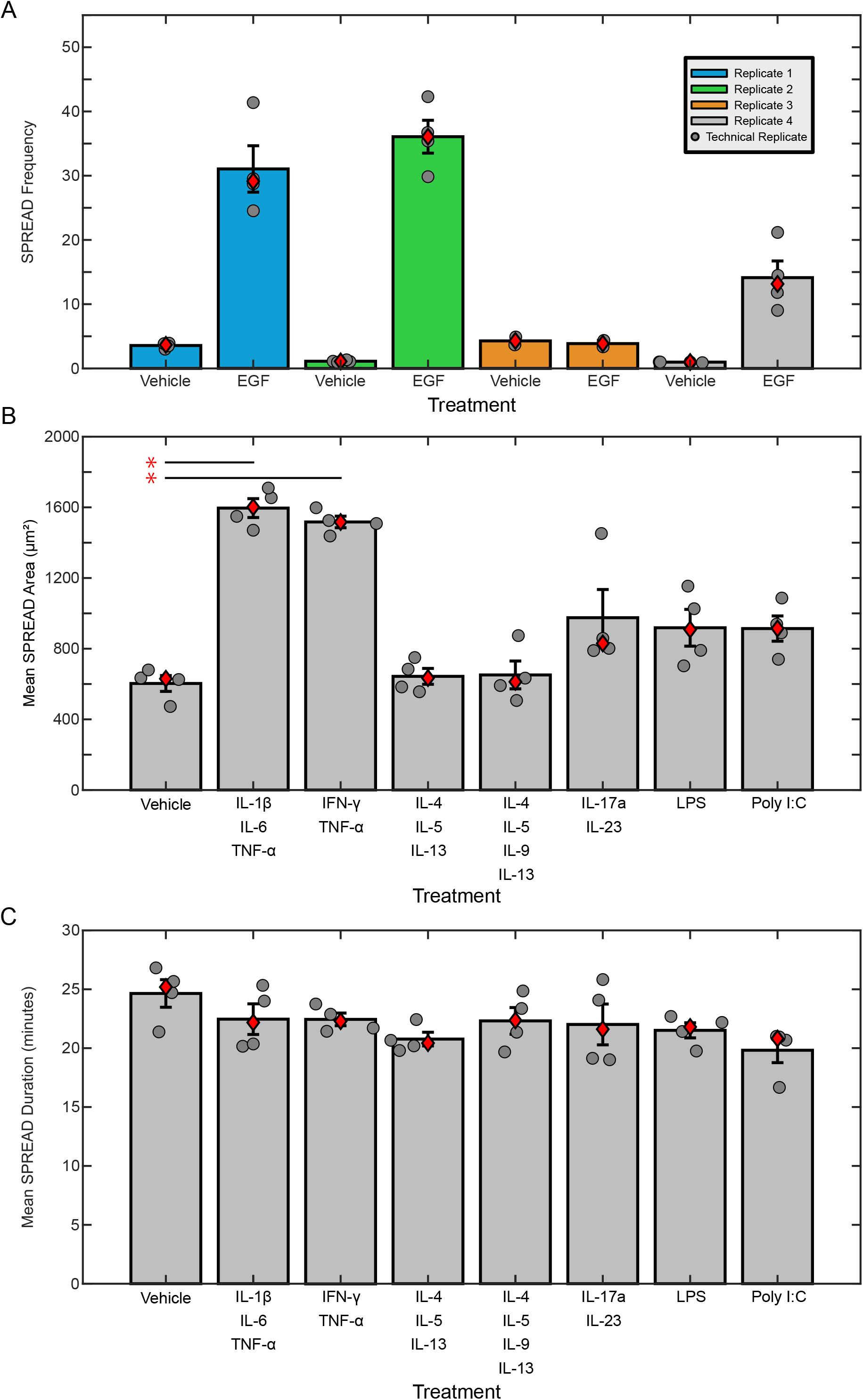
Variable SPREAD recognition in EGF treated cells and pro-inflammatory ligands-induced SPREAD size and duration in HBE1 cells. **(A)** SPREAD frequency in vehicle and EGF treated HBE1 cells compared across experimental replicates. Dark gray dots represent technical replicates within experimental replicates. **(B)** Mean SPREAD area (maximum area of per SPREAD in µm^2^) plotted according to treatment. **(C)** Mean SPREAD duration (in minutes) by treatment. For A–C, dark gray dots represent technical replicates, red diamonds indicate data median, and error bars show S.E.M. Red asterisks and lines indicate conditions where SPREAD duration is significantly different compared to vehicle control group (P < 0.005), calculated using 1-way ANOVA compared to the control and adjusted for multiple comparisons via the Dunnett’s procedure.

**Figure E4:**
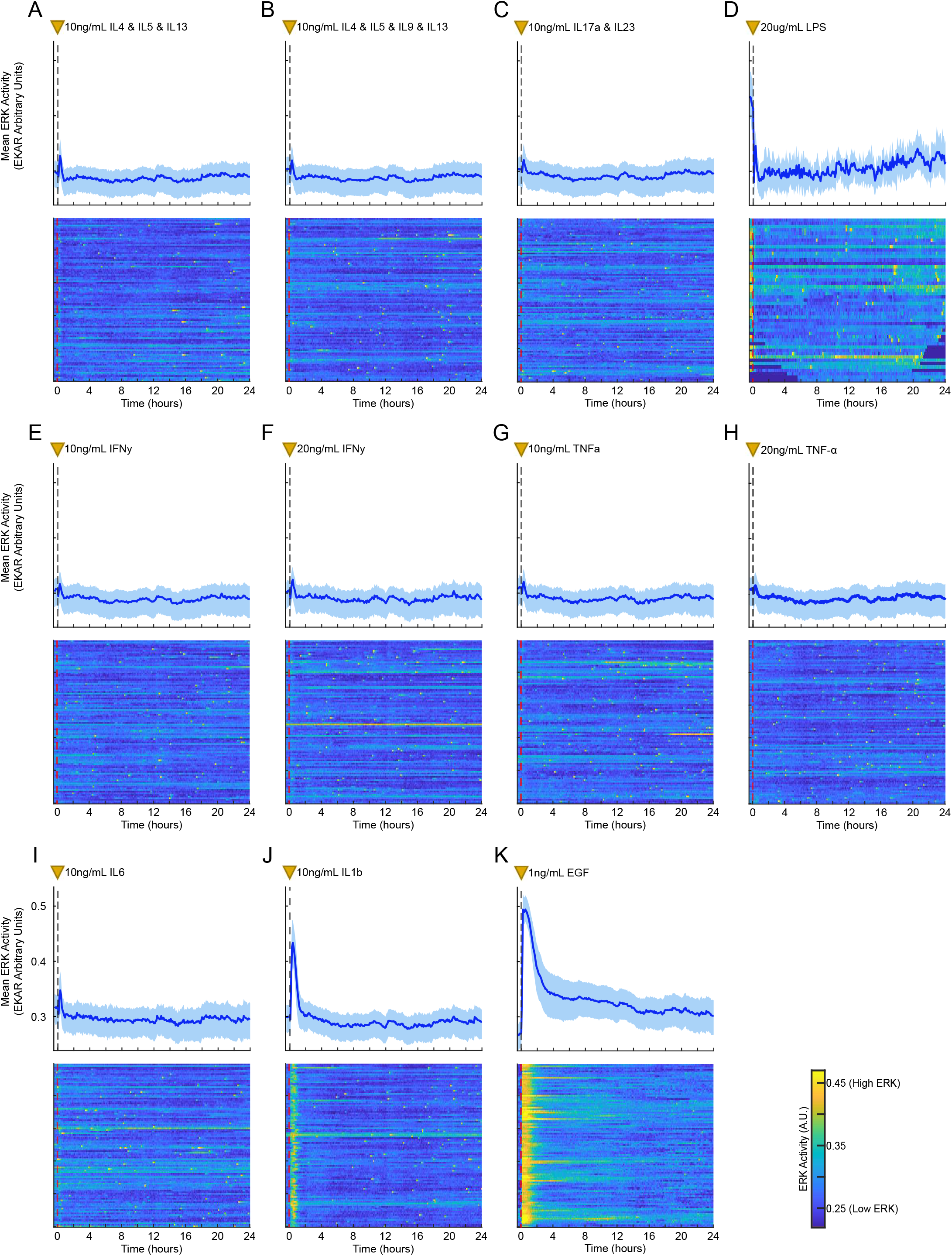
ERK responses of 16HBE cells to control and pro-inflammatory conditions. Top rows show mean with interquartile range of ERK activity across all cells from each condition. Heatmaps show ERK activity (color-axis) of 100 randomly selected cells from the data that make up the mean data plots above. Plots are representative data from 1 of 3 technical replicates, >500 cells per condition. Treatments occur at 0 hours, indicated by the dashed vertical lines and yellow triangles.

**Figure E5.**
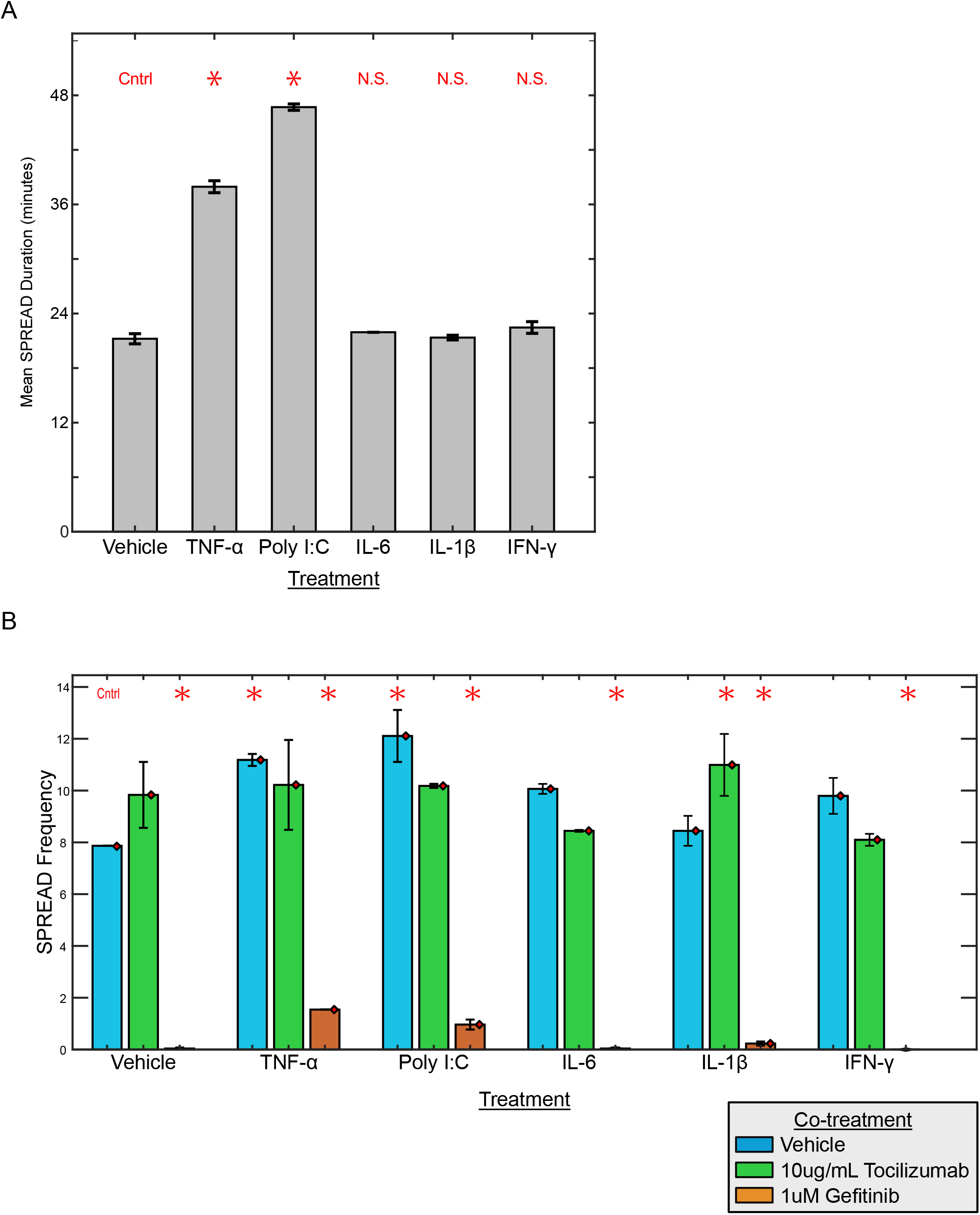
Receptor dependency of SPREADs in 16HBE cells. (A) Mean SPREAD duration (in minutes) by treatment. (B) SPREAD frequency of 16HBE cells co-treated with vehicle or pro-inflammatory conditions plus vehicle, tocilizumab, or gefitinib. Bars indicate the mean and red diamonds the median of the replicates. Error bars show S.E.M. Red asterisks indicate conditions where SPREAD frequency is significantly different compared to the control group (P < 0.005), calculated via 1-way ANOVA compared to the control and adjusted for multiple comparisons via Dunnett’s procedure.

**Figure E6.**
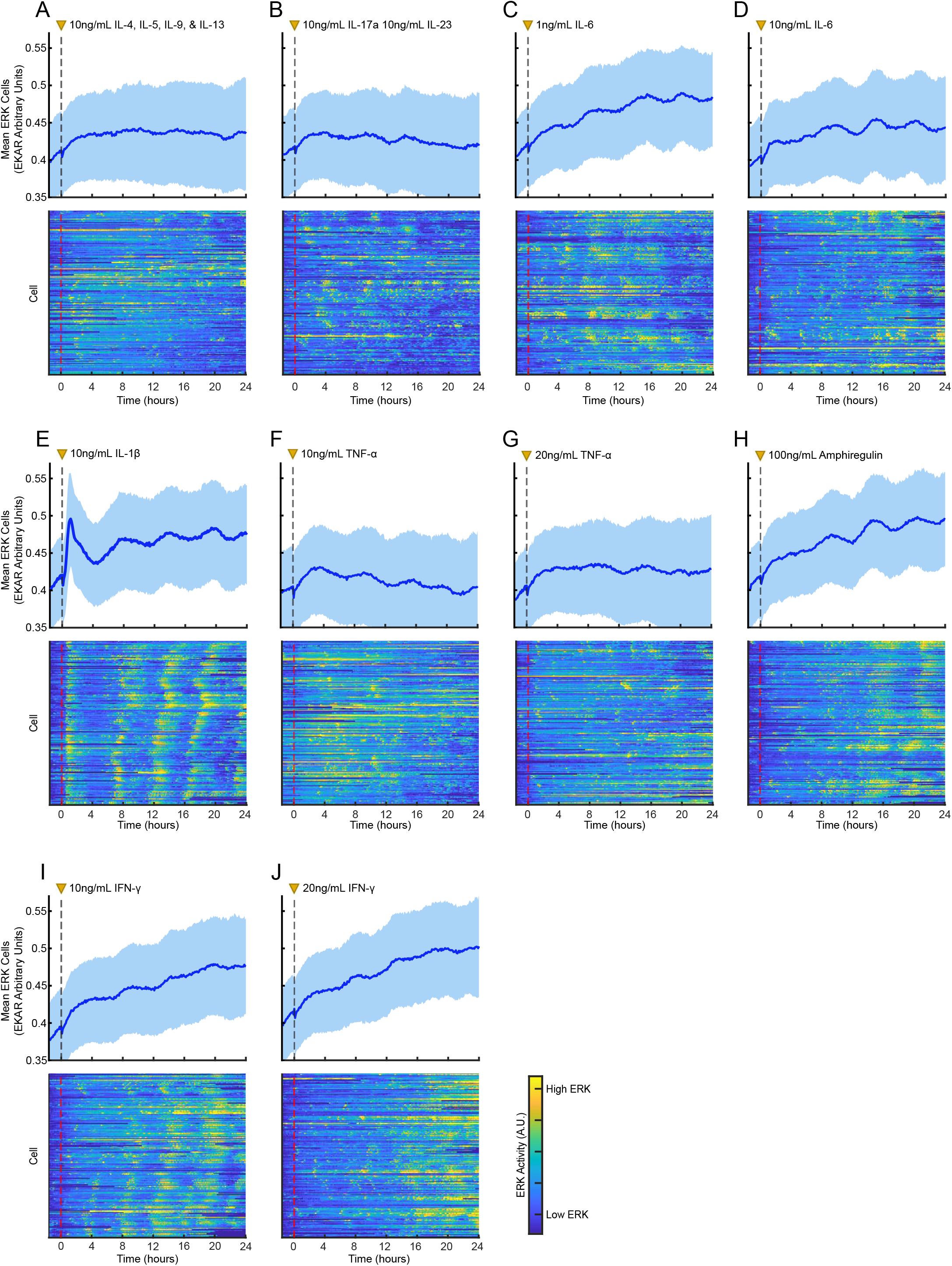
Mean and representative single cell ERK responses of pHBE* cells following treatment with pro-inflammatory conditions. (*Top rows*) Mean and 25^th^/75^th^ interquartile range of ERK data (EKAREN4) across all cells in representative experimental replicates (>4,000 cells per condition across 4 technical replicates). (*Bottom rows*) Heatmaps showing normalized ERK activity (color-axis) from 200 cells within a representative technical replicate, organized such that each cell (row) is positioned next to the closest neighboring cell in physical space. Treatment occurs at hour 0 as indicated by the dashed vertical line and yellow triangle. *primary human bronchial epithelial (pHBE) cells in submerged culture.

**Figure E7.**
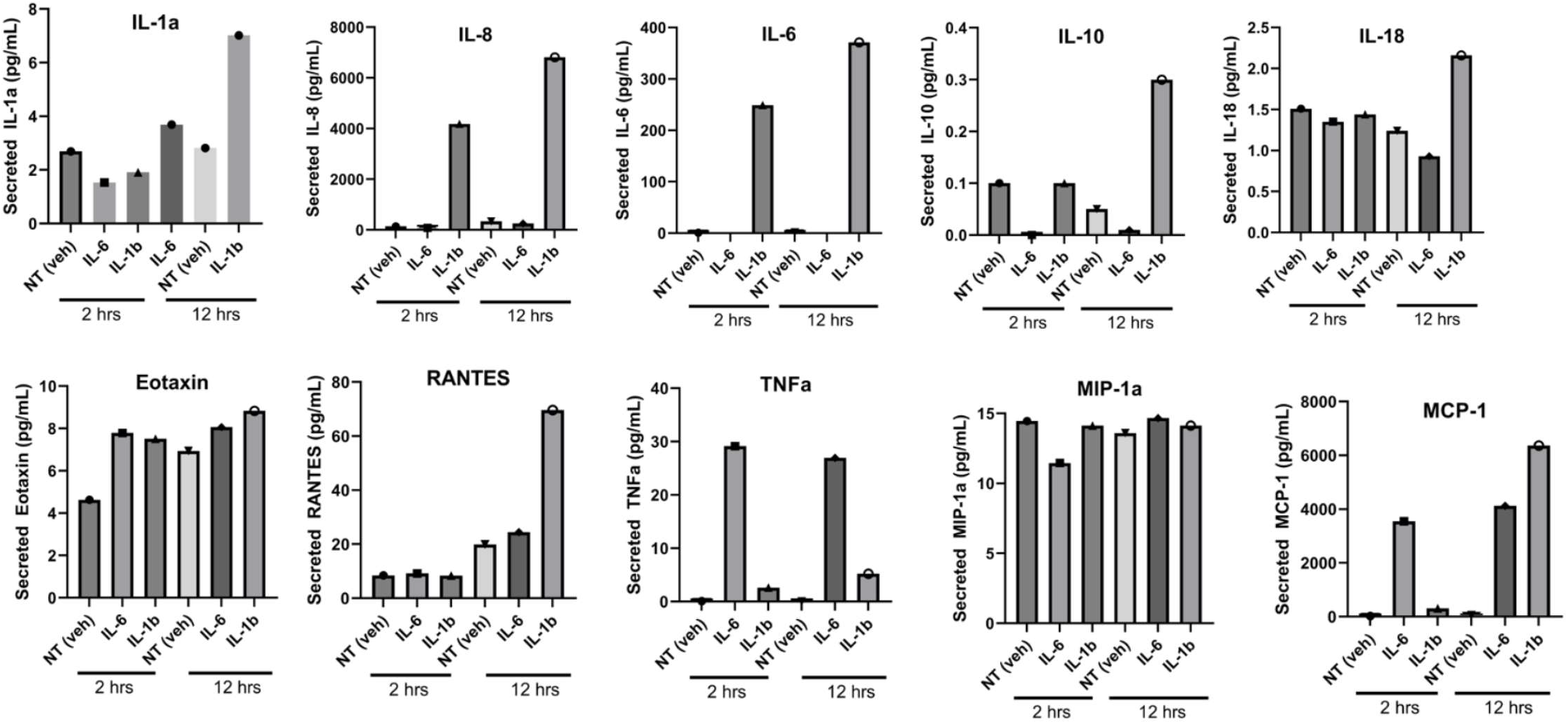
Luminex bead assay confirms cytokine release by HBE1 cells following IL-1β treatment. Bead-based cytokine assays for the indicated cytokines (plot titles) in cell culture supernatant from HBE1 cells treated with the cytokines listed on the horizontal axis.

**Figure E8.**
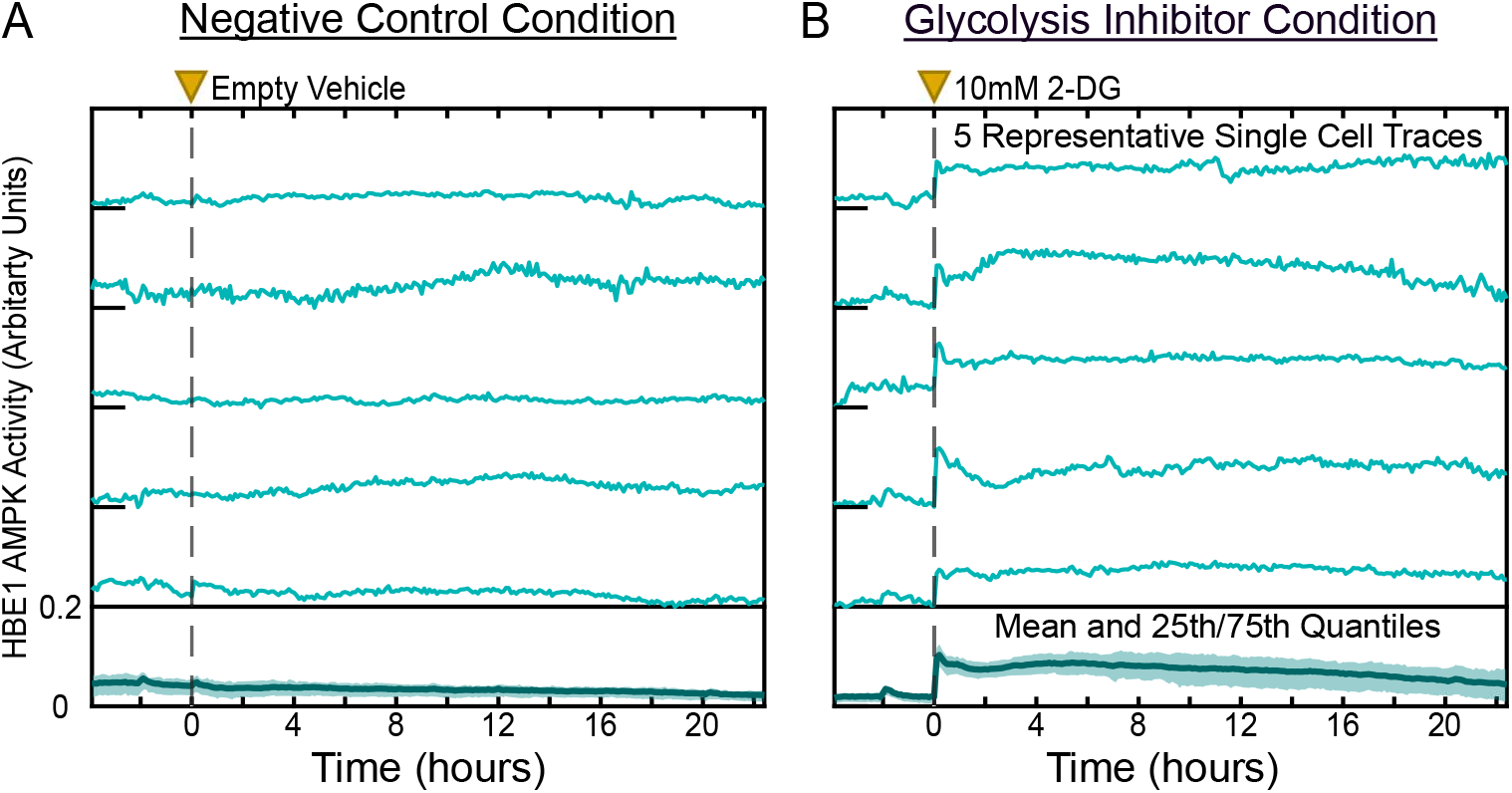
Verification of metabolic stress in HBE1 cells via monitoring of AMPK activity. Five representative single cell tracks and the mean (with 25^th^/75^th^ interquartile range) AMPK activity following treatment with **(A)** vehicle or **(B)** 10 mM 2-deoxyglucose (2-DG). Data are from 1 representative experimental replicate of 3 performed. Plots are derived from >500 cells of data per condition.

## SUPPLEMENTAL VIDEO LEGENDS

**Supplementary Video 1.** HBE1 cells that simultaneously express both ERK biosensors (EKAREN4 and ERK-KTR). *Top half* of the movie shows pseudo colored images made by taking the ratio of the CFP and YFP channels (EKAREN4). *Bottom half* of the movie shows ERK-KTR biosensor localization (red channel) in the same cells. Cells are treated with 10 ng/mL EGF near the conclusion of the movie. Single frames of this movie were used to make **Figure 1** of the paper.

**Supplementary Video 2.** SPREADs occurring amongst HBE1 cells expressing the ERK-KTR biosensor following treatment with 20 ng/mL IL-1β. Images from this experiment were used as images in **Figure 3A**.

**Supplementary Video 3.** Sporadic ERK activation across HBE1 cells following treatment with 10 ng/mL EGF. Pseudo colored movie showing EKAREN4 (ERK) activity; red coloration represents high ERK activity, while white coloration represents low ERK activity. Scale bar is 100 μm.

**Supplementary Video 4.** Untreated 16HBE cells showing SPREADs occurring after random events of apoptosis. Movie is pseudo colored showing the CFP/YFP ratio of EKAREN4 (ERK) activity. Scale bar is 250 μm.

**Supplementary Video 5.** pHBE* cells exhibiting wave-like ERK activation in response to 20 ng/mL IL-1β. Movie is pseudo colored by taking the ratio of the CFP and YFP channels, with red being high ERK activity and white being low ERK activity. *primary human bronchial epithelial (pHBE) cells in submerged culture. Scale bar is 260 μm.

**Supplementary Video 6.** Brightfield images of primary human bronchial epithelial cells after >30 days in air-liquid interface culture. Images show ciliary beating on the apical side of cells and was used to confirm differentiation prior to use in experiments.

**Supplementary Video 7.** ERK activation and SPREADs following treatment with 20 ng/mL IL-1β in differentiated primary human bronchial epithelial cells cultured in air-liquid interface. Pseudo colored images are the ratio of the CFP and YFP channels of the EKAREN4 biosensor; where red represents high ERK activity and white represents low ERK activity. Scale bar is 100 μm. Single frames from this movie are also used to make Figure 6A&B.

## SUPPLEMENTAL TABLES

Media Composition Tables:

**Table E1:** HBE1 cell culture medium.

| <u>Component</u> | <u>Vendor</u> | <u>Catalog Number</u> | <u>Final Concentration</u> |
| --- | --- | --- | --- |
| DMEM/F12 | GIBCO | 11320-033 | - |
| Cholera Toxin | Sigma | C8052 | 10 ng/mL |
| Endothelial Cell Growth Supplement from Bovine Neural Tissue Extract (BNE) | Sigma | E2759 | 7.5 ug/mL |
| Epidermal Growth Factor (EGF) | Peprotech | AF-100-15 | 10 ng/mL |
| Hydrocortisone | Sigma | H0888 | 0.5 ug/mL |
| Insulin | Sigma | I9278 | 5 ug/mL |
| Transferrin | Sigma | T2252 | 5 ug/mL |

**Table E2:** pHBE cell expansion medium.

| <u>Component</u> | <u>Vendor</u> | <u>Catalog Number</u> | <u>Final Concentration</u> |
| --- | --- | --- | --- |
| BEBM | Lonza | CC-3171 | - |
| Epidermal Growth Factor | Lonza | CC-4230F | 0.1% |
| Insulin | Lonza | CC-4021F | 0.1% |
| Transferrin | Lonza | CC-4205F | 0.1% |
| Hydrocortisone | Lonza | CC-4031F | 0.1% |
| Epinephrine | Lonza | CC-4221F | 0.1% |
| Triiodothyronine | Lonza | CC-4211F | 0.1% |
| Bovine Pituitary Extract | Lonza | CC-4009F | 0.4% |
| Penicillin | Gibco | 15140-122 | 15 Units/ml |
| Streptomycin | Gibco | 15140-122 | 15 ug/ml |
| Gentamycin | Lonza | CC-4081F | 0.1% |
| Amphotericin B | Gibco | 041-95780D | 0.25 mg/ml |
| Y-27632 (ROCK inhibitor) | Tocris Bioscience | 1254 | 5 uM |

**Table E3:** Primary human airway epithelial cell differentiation / ALI media. *All Trans Retinoic Acid was added after removal of apical growth medium from cells.

| <u>Component</u> | <u>Vendor</u> | <u>Catalog Number</u> | <u>Final Concentration</u> |
| --- | --- | --- | --- |
| DMEM/F12 +15mM HEPES | Corning | 10-092-CM | - |
| Epidermal Growth Factor | Millipore | GF-001 | 5 ng/ml |
| Insulin | Sigma | I-6634 | 4 ug/ml |
| Transferrin | Sigma | T-8158 | 5 ug/ml |
| Dexamethasone | Sigma | D-1756 | 100 nM |
| Cholera Toxin | List Biological Labs | 100B | 20 ng/ml |
| Bovine Hypothalamus Extract (BNE) | Lab Prepared | - | 0.2% |
| Bovine Serum Albumin | Sigma | A-9418 | 0.5 mg/ml |
| Ethanolamine | Sigma | E-0135 | 80 μM |
| MgCl<sub>2</sub> | Sigma | M-2393 | 0.3 mM |
| MgSO<sub>4</sub> | Sigma | M-2643 | 0.4 mM |
| Penicillin | Gibco | 15140-122 | 15 Units/ml |
| Streptomycin | Gibco | 15140-122 | 15 ug/ml |
| Gentamycin | Gibco | 15710-072 | 50 ug/ml |
| Amphotericin B | Gibco | 041-95780D | 0.25 mg/ml |
| All Trans Retinoic Acid* | Sigma | R-2625 | 30 nM |

**Table E4:** HBE1 imaging medium. *Insulin or Hydrocortisone were not added in experiments where insulin deprivation or hydrocortisone removal was tested.

| <u>Component</u> | <u>Vendor</u> | <u>Catalog Number</u> | <u>Final Concentration</u> |
| --- | --- | --- | --- |
| Base Imaging Medium - DMEM/F12 (equivalent to GIBCO 11320-033) with custom modifications; lacking glucose, glutamine, pyruvate, riboflavin, folic acid, and phenol red. | UC Davis Veterinary Medicine Biological Media Service | Custom | - |
| D-Glucose | Fisher | D16 | 17.4 mM |
| L-Glutamine | GIBCO | 35050079 | 2 mM |
| Sodium Pyruvate | GIBCO | 11360070 | 1 mM |
| Bovine Serum Albumin | Invitrogen | 16050-122 | 0.1% w/v |
| Cholera Toxin | Sigma | C8052 | 10 ng/mL |
| Hydrocortisone* | Sigma | H0888 | 0.5 ug/mL |
| Insulin* | Sigma | I9278 | 5 ug/mL |
| Transferrin | Sigma | T2252 | 5 ug/mL |

**Table E5:**
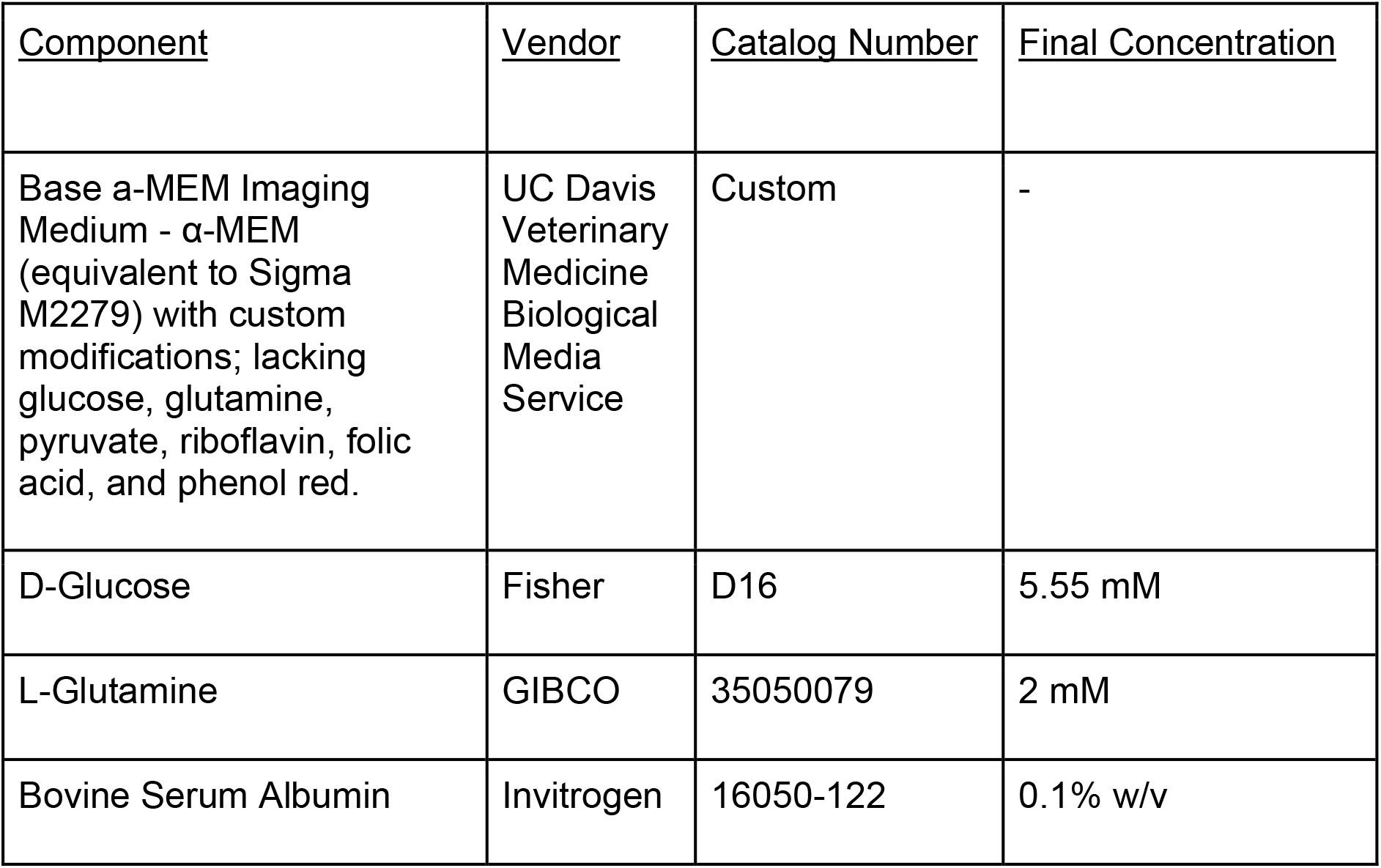
16HBE imaging medium.

| <u>Component</u> | <u>Vendor</u> | <u>Catalog Number</u> | <u>Final Concentration</u> |
| --- | --- | --- | --- |
| Base a-MEM Imaging Medium - α-MEM (equivalent to Sigma M2279) with custom modifications; lacking glucose, glutamine, pyruvate, riboflavin, folic acid, and phenol red. | UC Davis Veterinary Medicine Biological Media Service | Custom | - |
| D-Glucose | Fisher | D16 | 5.55 mM |
| L-Glutamine | GIBCO | 35050079 | 2 mM |
| Bovine Serum Albumin | Invitrogen | 16050-122 | 0.1% w/v |

**Table E6:**
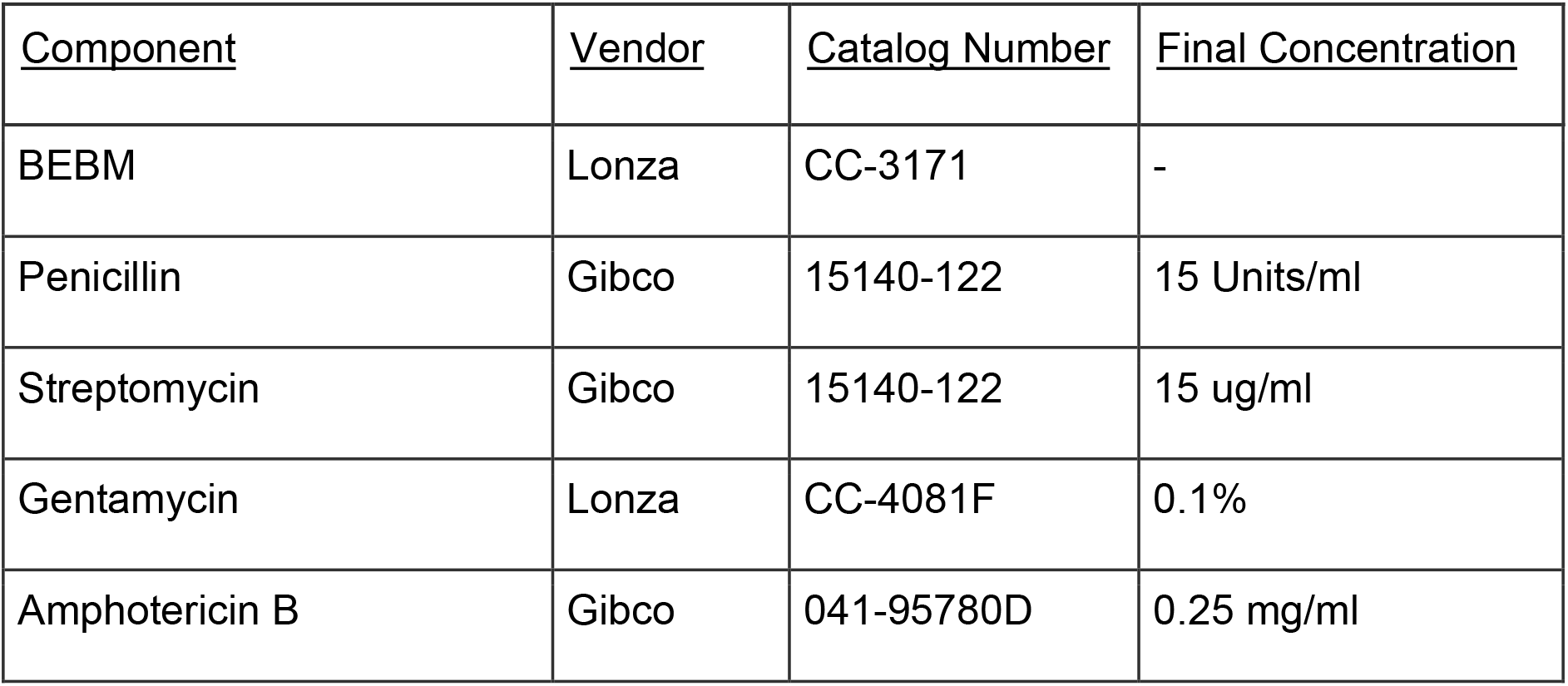
Submerged pHBE imaging medium.

| <u>Component</u> | <u>Vendor</u> | <u>Catalog Number</u> | <u>Final Concentration</u> |
| --- | --- | --- | --- |
| BEBM | Lonza | CC-3171 | - |
| Penicillin | Gibco | 15140-122 | 15 Units/ml |
| Streptomycin | Gibco | 15140-122 | 15 ug/ml |
| Gentamycin | Lonza | CC-4081F | 0.1% |
| Amphotericin B | Gibco | 041-95780D | 0.25 mg/ml |

**Table E7:** Air-Liquid Interface pHBE imaging medium.

| <u>Component</u> | <u>Vendor</u> | <u>Catalog Number</u> | <u>Final Concentration</u> |
| --- | --- | --- | --- |
| DMEM/F12 +15mM HEPES | Corning | 10-092-CM | - |
| Bovine Serum Albumin | Sigma | A-9418 | 0.5 mg/ml |
| Ethanolamine | Sigma | E-0135 | 80 μM |
| MgCl<sub>2</sub> | Sigma | M-2393 | 0.3 mM |
| MgSO<sub>4</sub> | Sigma | M-2643 | 0.4 mM |
| Penicillin | Gibco | 15140-122 | 15 Units/ml |
| Streptomycin | Gibco | 15140-122 | 15 ug/ml |
| Gentamycin | Gibco | 15710-072 | 50 ug/ml |
| Amphotericin B | Gibco | 041-95780D | 0.25 mg/ml |
| All Trans Retinoic Acid* | Sigma | R-2625 | 30 nM |

